# PGRMC1 phosphorylation and cell plasticity 2: genomic integrity and CpG methylation

**DOI:** 10.1101/737783

**Authors:** Bashar M. Thejer, Partho P. Adhikary, Sarah L. Teakel, Johnny Fang, Paul A. Weston, Saliya Gurusinghe, Ayad G. Anwer, Martin Gosnell, Jalal A. Jazayeri, Marina Ludescher, Lesley-Ann Gray, Michael Pawlak, Robyn H. Wallace, Sameer D. Pant, Marie Wong, Tamas Fischer, Elizabeth J. New, Tanja N. Fehm, Hans Neubauer, Ewa M. Goldys, Jane C. Quinn, Leslie A. Weston, Michael A. Cahill

## Abstract

Progesterone receptor membrane component 1 (PGRMC1) is often elevated in cancers, and exists in alternative states of phosphorylation. A motif centered on PGRMC1 Y180 was evolutionarily acquired concurrently with the embryological gastrulation organizer that orchestrates vertebrate tissue differentiation. Here, we show that mutagenic manipulation of PGRMC1 phosphorylation alters cell metabolism, genomic stability, and CpG methylation. Each of several mutants elicited distinct patterns of genomic CpG methylation. Mutation of S57A/Y180/S181A led to increased net hypermethylation, reminiscent of embryonic stem cells. Pathways enrichment analysis suggested modulation of processes related to animal cell differentiation status and tissue identity, as well as cell cycle control and ATM/ATR DNA damage repair regulation. We detected different genomic mutation rates in culture. A companion manuscript shows that these cell states dramatically affect protein abundances, cell and mitochondrial morphology, and glycolytic metabolism. We propose that PGRMC1 phosphorylation status modulates cellular plasticity mechanisms relevant to early embryological tissue differentiation.

## INTRODUCTION

Progesterone (P4) Receptor Membrane Components 1 (PGRMC1) is a cytochrome b_5_ (cytb5) protein and a member of the membrane-associated P4 receptor (MAPR) family with a plethora of cellular functions. These include some cytb5-typical functions that predate or are potentially ancient in eukaryotes (regulation of heme synthesis (Piel et al., 2016), cyP450 (cytochrome P450) interactions (Oda et al., 2011; Ryu et al., 2017; Szczesna-Skorupa and Kemper, 2011), sterol metabolism (Cahill and Medlock, 2017; Hughes et al., 2007; Mallory et al., 2005)), some that evidently arose in eukaryotes (membrane trafficking (reviewed by: Cahill et al., 2016a; Mifsud and Bateman, 2002), cell cycle regulation at the G0/G1 checkpoint (Cahill et al., 2016a; Griffin et al., 2014; Peluso et al., 2014), mitotic/meiotic spindle association (Juhlen et al., 2018; Luciano and Peluso, 2016; Terzaghi et al., 2016)), and clearly specialized metazoan functions including e.g., fertility, embryogenic axon guidance, and membrane trafficking associated with synaptic plasticity (reviewed by: Cahill et al., 2016a; Rohe et al., 2009).

CyP450-interactions (Ryu et al., 2017) conspicuously feature PGRMC1 regulation of the most conserved eukaryotic cyP450 (lanosterol 14-alpha demethylase, CYP51A) to modify the first sterol (lanosterol) from yeast to mammals (Cahill, 2007; Hughes et al., 2007; Mallory et al., 2005). The protein is accordingly found in diverse subcellular locations, including endoplasmic reticulum/perinuclear membrane and Golgi apparatus (Neubauer et al., 2008; Sakamoto et al., 2004), at the cell surface (Izzo et al., 2014; Kim et al., 2019), mitochondria (Piel et al., 2016), and the nucleus, where it localizes to nucleoli (Terzaghi et al., 2018). It is strongly associated with sterol biology (reviewed by: (Cahill, 2007; Cahill and Medlock, 2017)), and a complex between PGRMC1, the low density lipoprotein (LDL) receptor (LDLR) and the sigma-2 receptor transmembrane protein 97 (TMEM97) regulates the rapid internalization of LDLR (Riad et al., 2018).

Indeed, PGRMC1-associated membrane trafficking determines the cell surface localization of a number of different proteins including e.g., epidermal growth factor receptor (Ahmed et al., 2010a), insulin receptor and glucose transporters GLUT-1 and GLUT-4 in A549 lung cancer cells (Hampton et al., 2018), glucagon-like peptide-1 (GLP-1) receptor complex in pancreatic beta cells (Zhang et al., 2014), synaptic vesicles in the central nervous system (Izzo et al., 2014), and membrane progestin receptor α (mPRα) in rat granulosa cells (Thomas et al., 2014) and maturing zebrafish oocytes (Aizen et al., 2018). The translocation of transmembrane proteins such as membrane receptors to the cell surface could underlie much of the biology attributed to PGRMC1 (Ahmed et al., 2010b).

Anti-apoptotic activity of PGRMC1, especially to treatment with the topoisomerase II inhibitor doxorubicin, has been reported by several studies (Friel et al., 2015; Lin et al., 2015). This activity was linked to P4- and PGRMC1-dependent survival responses (Peluso et al., 2008). It remains unclear how much this P4-responsiveness involves the membrane trafficking of actual P4 receptors to the cell surface, or to what extent PGRMC1 itself may mediate its own P4 response (reviewed by (Cahill et al., 2016a; Rohe et al., 2009)). PGRMC1 undeniably confers P4 responsiveness to cells of the reproductive (Peluso and Pru, 2014) and the central nervous systems (Petersen et al., 2013). A D120G heme-binding incapacitated mutant also exhibited reduced P4-responsiveness, suggesting independence from cyP450 activation (Peluso et al., 2008). PGRMC1 associates with the CYP2D6 and CYP3A4 cyP450 enzymes responsible for doxorubicin hydroxylation and inactivation, and this activity required the heme-chelating residue Y113 (Kabe et al., 2016). However, it remains unclear whether heme-chelation or tyrosine phosphorylation (and perhaps membrane trafficking) of Y113 is involved (Cahill et al., 2016b).

PGRMC1 is induced by a variety of agents causing DNA damage, including doxorubicin (Rohe et al., 2009), and its yeast Dap1 (*<u>d</u>*amage-*<u>a</u>*ssociated *<u>p</u>*rotein 1) homologs are associated with DNA damage response (Hand et al., 2003; Hughes et al., 2007). Heme-binding and cyP450 interactions are thought to be required for DNA damage response (Hughes et al., 2007; Rohe et al., 2009).

A549 cells treated with so-called PGRMC1 inhibitor AG-205 exhibited G1 arrest and an increase in sub-G1 (apoptotic) cells, with a reduction in the S and G2/M stages (Ahmed et al., 2010b). Note that AG-205 is often cited as a PGRMC1-specific inhibitor. While it certainly affects PGRMC1-dependent function, it was developed against a plant MAPR protein (Yoshitani et al., 2005). There is no evidence that it is any more specific to PGRMC1 than to PGRMC2, other MAPR family members Neudesin or Neuferricin, or indeed that it does not exert effects involving non MAPR proteins. Claims related to “PGRMC1-specific inhibitor AG-205” should be treated circumspectly.

The PI3K/Akt pathway is affected by PGRMC1 in MCF-7 cells by PGRMC1 overexpression (Hand and Craven, 2003) or overexpression of phosphorylation site mutants (Neubauer et al., 2008). In an accompanying paper we show that Y180 is required for PI3K/Akt activity (Thejer et al., 2019). PGRMC1 was differentially phosphorylated between estrogen receptor-positive and negative breast cancers (Neubauer et al., 2008). Human PGRMC1 contains predicted binding site motifs for Src homology 2 (SH2) and Src homology 3 (SH3) domain-containing proteins, with several other phosphorylation sites at S57, T178 and S181 being thought to additionally regulate these sites (Cahill et al., 2016b). S57 and the SH2 target motifs containing Y139 and T178/Y180/S181 are mutually juxtaposed on the folded protein surface, forming a potential proximity-stimulated signaling platform (Cahill, 2007; Cahill et al., 2016b). Thus, PGRMC1 represents a potential integration point and effector of many cell signaling pathways responsible for growth and proliferation (Ahmed et al., 2010b; Cahill, 2007; Cahill et al., 2016a; Peluso and Pru, 2014).

In an accompanying paper (Thejer et al., 2019), we found that mutating hemagglutinin (HA)-tagged PGRMC1 (PGRMC1-HA) S57 and S181 induced dramatic effects in MIA PaCa-2 (MP) pancreatic cancer cells. This double mutant (DM) exhibited elevated PI3K/Akt activity, altered actin cytoskeleton proteins, and elevated migration rates relative to MP cells expressing wild-type (WT) PGRMC1-HA protein. Additional mutation of Y180 in a triple mutant (TM) attenuated PI3K/Akt activity and mouse xenograft tumor growth. Pathways enrichment analysis of a proteomics study suggested broad effects relevant to cancer biology, including effects on glucose metabolism and mitochondrial form and function that were suggestive of plastic cellular reprogramming. Here, we characterize those cells further considering effects on metabolism, genomic stability and epigenetic plasticity.

## RESULTS

### PGRMC1 phosphorylation state-dependent metabolic differences

Cells were subjected to preliminary hyperspectral autofluorescence imaging: a non-invasive analytical technique (Elliott et al., 2012). Distinct spectral signatures of endogenous auto-fluorescent molecules (Gosnell et al., 2016b; Lu and Fei, 2014) can provide valuable information about intracellular metabolic state (Gosnell et al., 2016a). As a first step towards demonstrating discrimination of cell conditions based on spectral information, eight informative spectral features that discriminated the cells were chosen (Figure 1A and Table S1). The cells in each condition could be clearly distinguished using the autofluorescence of their endogenous metabolites. A pairwise linear discrimination analysis using the same spectral features indicated that all cell conditions were highly significantly separated from each other on the basis of endogenous autofluorescence (Figure 1A).

**Figure 1.**
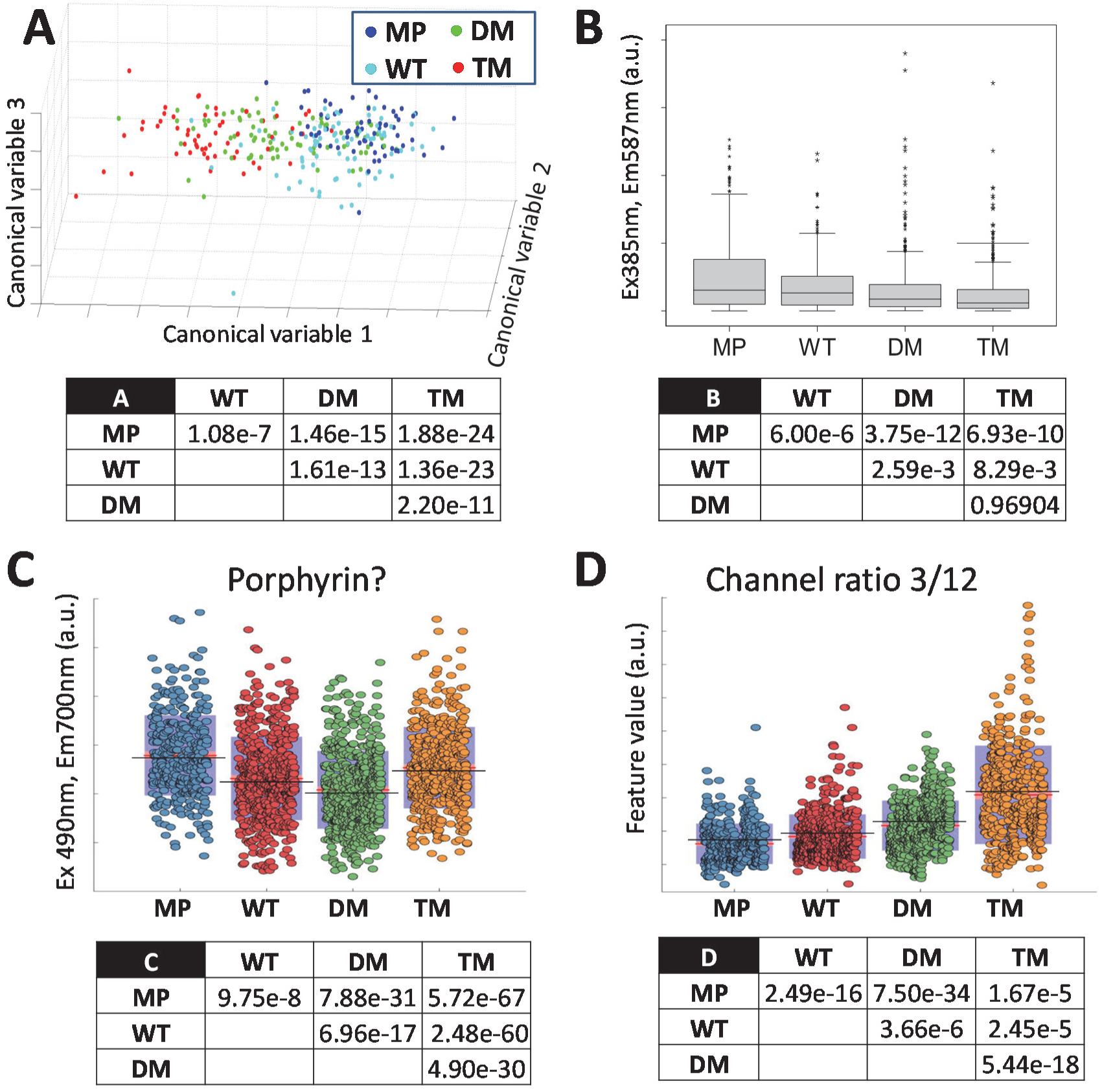
Hyperspectral autofluorescence reveals metabolic differences between cells expressing different mutant PGRMC1-HA proteins. (A) Principal component analysis (PCA) followed by linear discriminant analysis (LDA) reveal hyperspectral autofluorescence parameters that significantly discriminate between cell types. The panel provides a three dimensional depiction of the three most significant canonical variables revealed by PCA and LDA as described (Gosnell et al., 2016a) that differ according to PGRMC1-HA phosphorylation status. The table provides *p* values from Kolmogorov-Smirnov test pairwise discrimination analysis. (B) Mean cellular intensity of hyperspectral autofluorescence channel 3 [375nm(Ex), 450nm(Em)], which corresponds primarily to flavin emission, is significantly affected by PGRMC1-HA phosphorylation status. Boxplots were generated in SPSS. The table shows Kruskal-Wallis test *p* values for the pair-wise condition comparisons performed on primary emission data.

Figure 1B shows autofluorescence differences in spectral channel 3, thought to reflect flavin emission (Gosnell et al., 2016a; Huang et al., 2002). All cells over-expressing PGRMC1-HA proteins exhibited reduced levels, with DM and TM the lowest (Figure 1B). We also noted significant differences observed between cell lines for autofluorescent emission at 700 nm (Figure S1A), in a wavelength range at which heme-containing proteins and other porphyrins contribute strongly to autofluorescence (Schneckenburger et al., 1989). Note that PGRMC1 not only binds heme but is involved in heme synthesis (Piel et al., 2016). Heme itself does not emit fluorescence but leads to fluorescence quenching, however, various heme-containing proteins are fluorescent (Schneckenburger et al., 1989). Although the actual identities of these and fluorescent species contributing to many hyperspectral channels remain unknown, several unidentified parameters also significantly discriminated between the metabolites present in the different cell lines, such as e.g. the ratio of channel 3 to channel 12 (Figure S1B,C, Table S1). These results indicated that PGRMC1 phosphorylation status dramatically alters cell metabolic state.

### PGRMC1 phosphorylation mutants did not affect P4 or AG-205 responses

We were interested in whether DNA mutation rates may have been related to the P4-dependent protection against cell death, or to the mechanism of AG-205-indiced cell death. Consistent with previously reported PGRMC1-dependent anti-mitotic effects of P4 in Ishikawa endometrial cancer cells (Friel et al., 2015) and MDA-MB-231 breast cancer cells (Clark et al., 2016), incubation of cells in 1μM P4 retarded cell proliferation of all cells over-expressing a PGRMC1-HA protein (WT, DM or TM), but not of MP cells relative to non-P4-treated control cells (Figure 3A). When cells pretreated with or without P4 for 1 hr were co-incubated in the presence doxorubicin (Dox), in the absence of P4 then WT, DM and TM cells were more susceptible to Dox-induced death (Figure 3B-C, white data points and boxes), however these cells exhibited greater P4-dependent protection against Dox-induced death (Figure 3B-C, shaded data points and boxes). As reported previously for Ishikawa cells (Friel et al., 2015), the 1hr pre-incubation with P4 was required to observe these protective effects of P4 (data not shown). There was no significant protective effect of P4 on cell survival for MP cells (Figure 3C). Notably, there were no significant differences in P4-dependent protection between WT, MP or TM cells. There was also no difference in the sensitivity of any of these cells to AG-205-induced cell death, with essentially superimposable dose response curves, and a half maximal lethal effective concentration (EC_50_) of approximately 32μM AG-205 (Figure S2). Therefore, although we observe PGRMC1-HA-dependent response to P4, this response is apparently unaffected by the influence of our PGRMC1 phosphorylation mutants. We conclude that the dramatic effects of altered PGRMC1 phosphorylation status observed in this and the accompanying manuscript (Thejer et al., 2019) are due to different and newly described functions of PGRMC1, unrelated to the mechanism of P4-induced vitality or AG-205-induced death.

### PGRMC1 phosphorylation status affects cytoplasmic redox status

Naphthalimide-flavin redox sensor 1 (NpFR1) is a fluorophore analogous to NpFR2 (Kaur et al., 2015; Thejer et al., 2019), except that it is localized to the cytoplasm (Yeow et al., 2014). To further explore metabolic state, cells were treated with NpFR1 and fluorescent state was assayed by flow cytometry to reveal two cell populations: one with less oxidized NpFR1 and one with more oxidized NpFR1 (Figure 2A). The bimodal fluorescent peaks of Figure 2A represent cell populations with different levels of cytoplasmic oxidation/reduction which have not been further characterized. All PGRMC1-HA-expressing cells exhibited the majority of cells in a more oxidized peak. TM (9.4%) and WT (17.7%) cells both exhibited less cells in the lower oxidized fraction (Figure 2A,B, black arrows). For TM cells, which had most cells in the more oxidizing cytoplasmic population, the degree of NpFR1 oxidation in that fraction was significantly lower than for all other cells (Figure 2A,C, white arrow). Higher cytoplasmic oxidative levels in TM cells suggested potential effects on genomic mutation rates.

**Figure 2.**
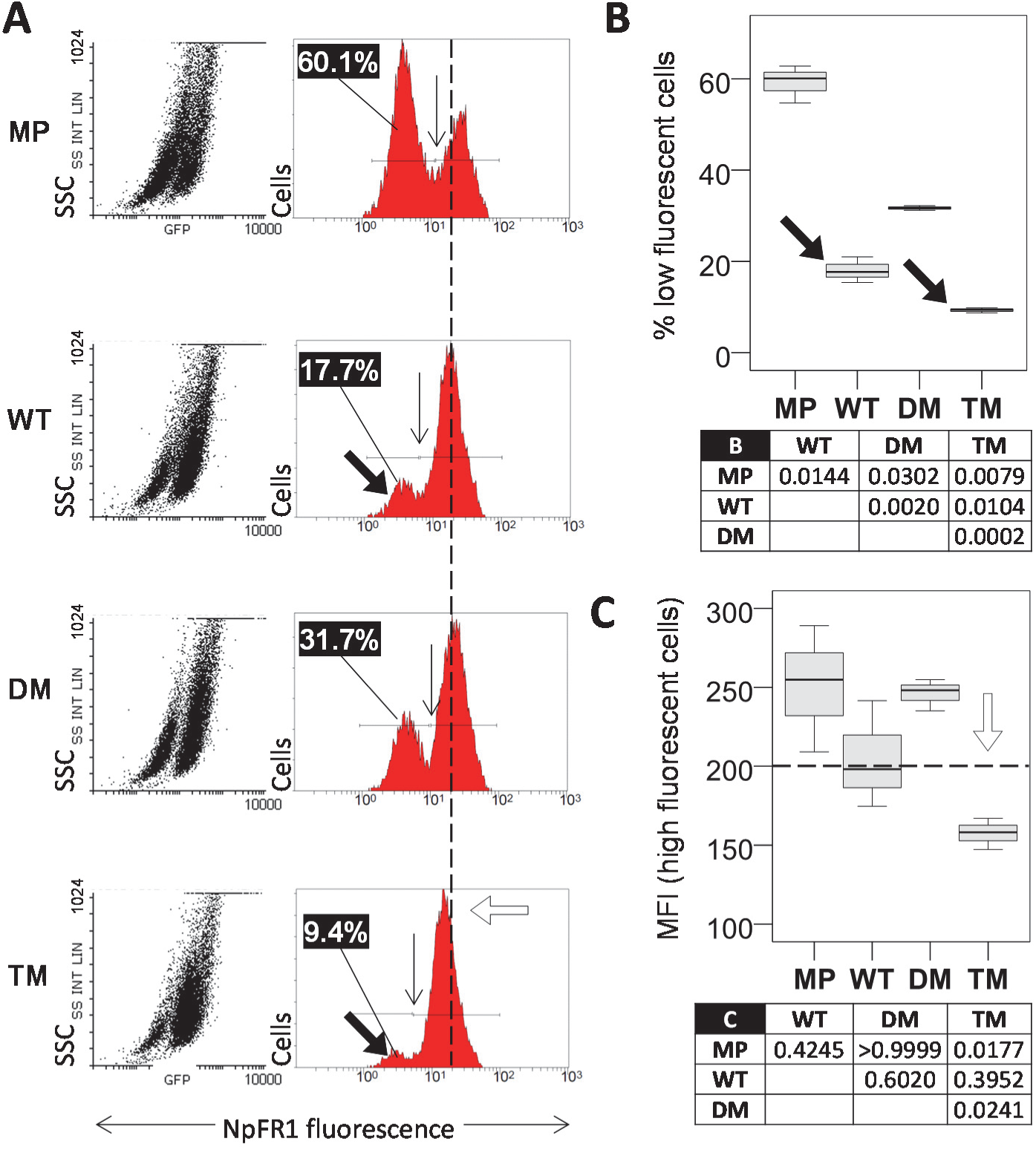
NpFR1 reveals differences in cytoplasmic redox status. (A) NpFR1 flow cytometry results. Scatter plots to the left depict green fluorescence on the x axis, and side scatter on the y axis. In the right panels the y axis represents cell number. The numbers at the upper left of each right panel are the fraction of the population in the respective left hand low fluorescent peak. Vertical arrows show the boundary between cell populations, and the inset numbers in the left of each panel represent the percentage of cells in the less oxidized left hand cell population. The vertical dotted reference line represents the 200 fluorescence intensity units, and the open arrow highlights TM fluorescence. (B) Boxplots showing mean and sample distribution for % cells in the less oxidized population of replicates of (A). Sample size is n=3, being one each of independent cell lines 1-3 for WT, DM and TM, and triplicates of the MP cell line. The table shows ANOVA post-hoc Dunnet’s T3 *p* values for the indicated pairwise comparisons. (C) Boxplot of median fluorescent intensity (MFI) of the oxidized cell populations from (A). The horizontal dotted reference line and open arrow are identical to (A). The table shows ANOVA post-hoc Bonferroni *p* values for the indicated pairwise comparisons.

**Figure 3.**
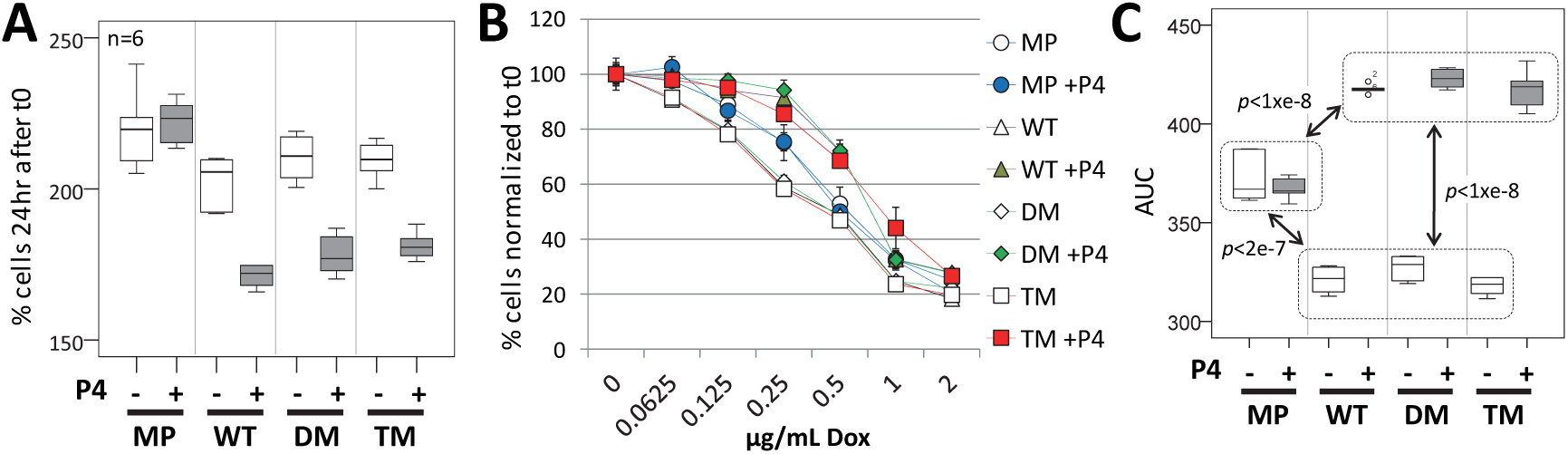
PGRMC1 phosphorylation status does not affect P4-dependent resistance to doxorubicin toxicity or resistance to AG-205-induced cell death. (A) P4 reduces cell proliferation of cells expressing all PGRMC1-HA proteins. The panel shows boxplots of viable cells for n=6 replicates. The viability of cells pretreated with P4 (dark boxes) or DMSO vehicle control (light boxes) after 23 hr were allowed to grow a further 24 hours and the level of MTT formazan was quantified as a proxy for viable cell numbers. “% cells after 48 hr” is presented relative to the signal obtained after adherence for 3 hr. No significant differences were observed between non-P4 treated cell pairings, except MP v. TM (*p*=0.021, 2 way ANOVA). Pair wise comparisons of P4-treated MP vs any of P4-treated WT, DM or TM, or of +/-P4 treatment for WT, DM and TM, were significantly different at the *p*<0.00001 level (2 way ANOVA). MP cells +/- P4 were not different (*p*=0.55, two way ANOVA). Considering only P4-treated cells, MP differed in P4 response from all other cell types (*p*<1×10^-10^), and WT differed marginally from TM (*p*=0.034) by one way ANOVA and post-hoc Bonferroni. (B) P4-protection of MP cells from doxorubicin-induced cell death is facilitated by over-expression of PGRMC1-HA proteins (WT, DM & TM). Cells were grown as in (A), except at t0 doxorubicin (Dox) was added at the indicated concentrations, followed by 24hr incubation. Because of altered cell proliferation during pretreatment with P4 (A), all signals at t0 were expressed as a percentage of the averaged control sample without Dox at t0 to assess the effects of P4. Data points represent the averages ± s.d. of n=6 replicates. (C) Boxplots of area under the curve (AUC) results for all data points with dox addition from (B). Two way ANOVA dox treatments were statistically significant (F=1292.237, df=1, df2=40, *p*< 1=e-8), with Partial Eta Squared indicating 97% effect size in the data. Considering pairwise comparisons of cell classes, DM v. TM (*p*=0.014) and WT v. DM (*p*=0.05) were significantly different. Pairwise comparison between the AUC levels for +/-P4 for each cell class yielded no difference for MP, but *p*<1×10^-8^ for other cell types, +/- P4. One way ANOVA post-hoc Bonferroni pairwise comparisons for -P4 cells revealed that samples within dotted boxes did not differ significantly, whereas all pairwise *p* values were less than those indicated for comparisons between boxes.

### PGRMC1 phosphorylation status affects pathways associated with replication, cell cycle, and mitogenic signaling

In an accompanying paper (Thejer et al., 2019) we presented only highly significant differential WebGestalt pathways with Benjamini-Hochberg adjusted *p* (adjP) values <0.001. Several pathways related to cell cycle control and DNA replication were also detected, albeit with weaker significance, including five proteins predicted to be associated with decreased levels of ATM/ATR DNA damage sensing and control in TM cells (Figure 4A), which was interesting to us because of the DNA damage associations of PGRMC1 (see Introduction). Although we could not detect differences in cultured proliferation of any cell lines (Thejer et al., 2019), reverse phase protein array (RPPA) analysis revealed elevated levels of activated MKK4, MEK and ERK, as well as phosphorylated retinoblastoma protein (Rb) in TM cells (Figure 4B). This suggests that PGRMC1 Y180 is somehow involved in regulating the G1 checkpoint, which may be related to the reported ability of PGRMC1 and/or PGRMC1 to lower the propensity of cells to enter the G1 and the cell cycle, rather than enter G0, in spontaneously immortalized granulosa cells (Peluso et al., 2014).

**Figure 4.**
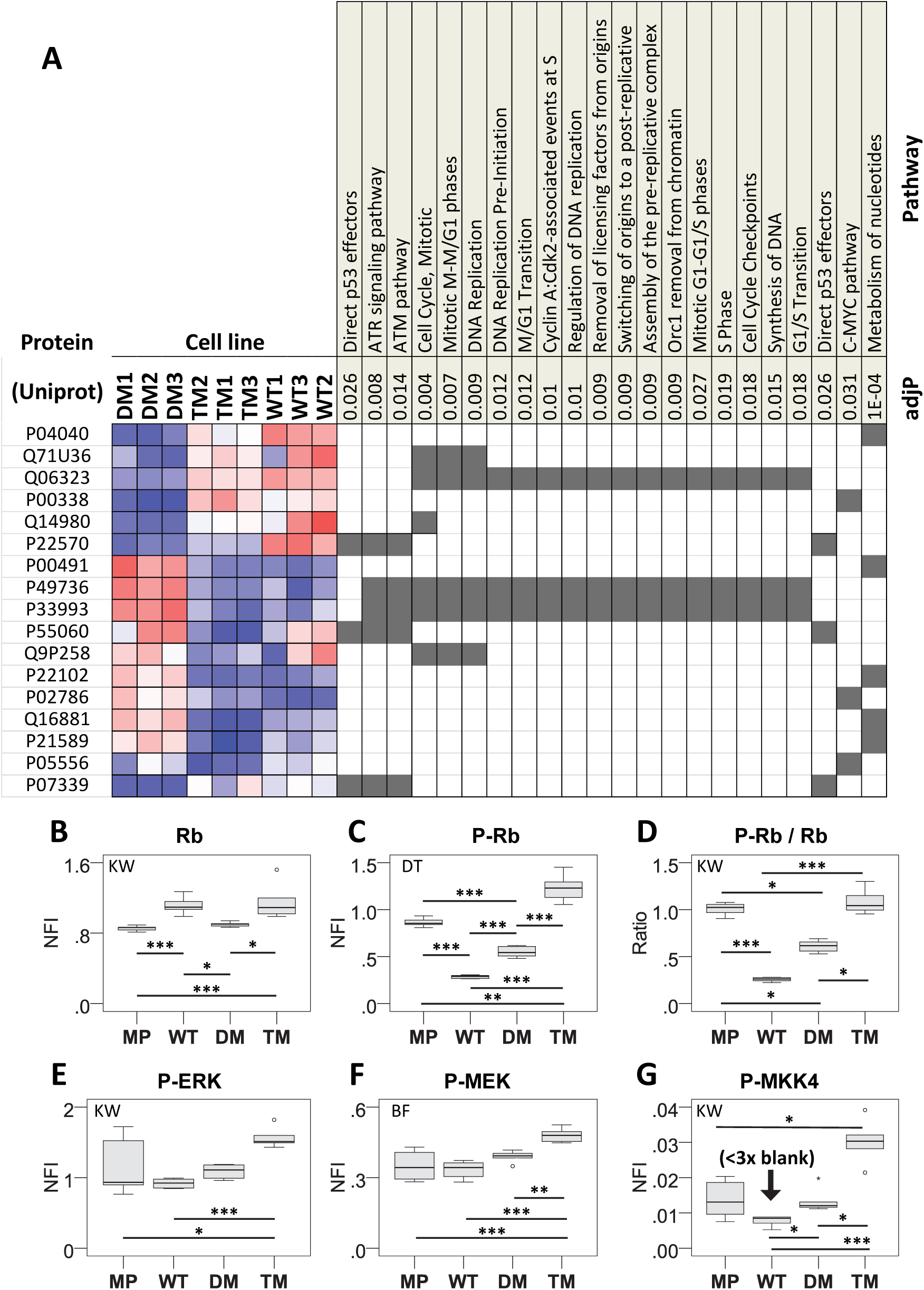
Pathways enrichment suggests altered cell cycle control proteins. (A) Abundances of proteins detected as significantly differentially abundant in indicated Pathways Commons pathways by enrichment analysis (Accompanying paper (Thejer et al., 2019)) at the adjP < 5% significance level. (B, C, E-G) Average reverse phase protein array (RPPA) normalized fluorescent intensity (NFI) from the indicated antibodies (see methods) is plotted from 6 replicate measurements. NFI is normalized to protein content. Significance levels are * *p*<0.05, ** *p*<0.01, *** *p*<0.001. Conventions follow those of the accompanying paper (Thejer et al., 2019). (D) The ratio of average NFI of the anti-pRB antibody to that of the RB antibody.

### PGRMC1 phosphorylation status affects genomic integrity

Altered cytoplasmic oxidative environment, potential perturbations of the ATM/ATR pathways, and G1 checkpoint features without detectable differences in proliferation rates (Thejer et al., 2019) led us to explore genomic stability. We cultured one cell line each of MP, WT, DM and TM for 30 passages, and sequenced the genomes of each. The PGRMC1-HA constructs were expressed at similar levels after 30 passages (Figure 5A). Relative to the reference human genome sequence, the order of average mutation rates (difference to reference) per chromosome were WT<DM<TM (not shown), which was also reflected in unique mutations per MB, as rates per chromosome (Figure 5B). This indicates that manipulation of the phosphorylation status of PGRMC1 Y180 can affect genomic stability, which is of potentially great relevance to cancer biology.

**Figure 5.**
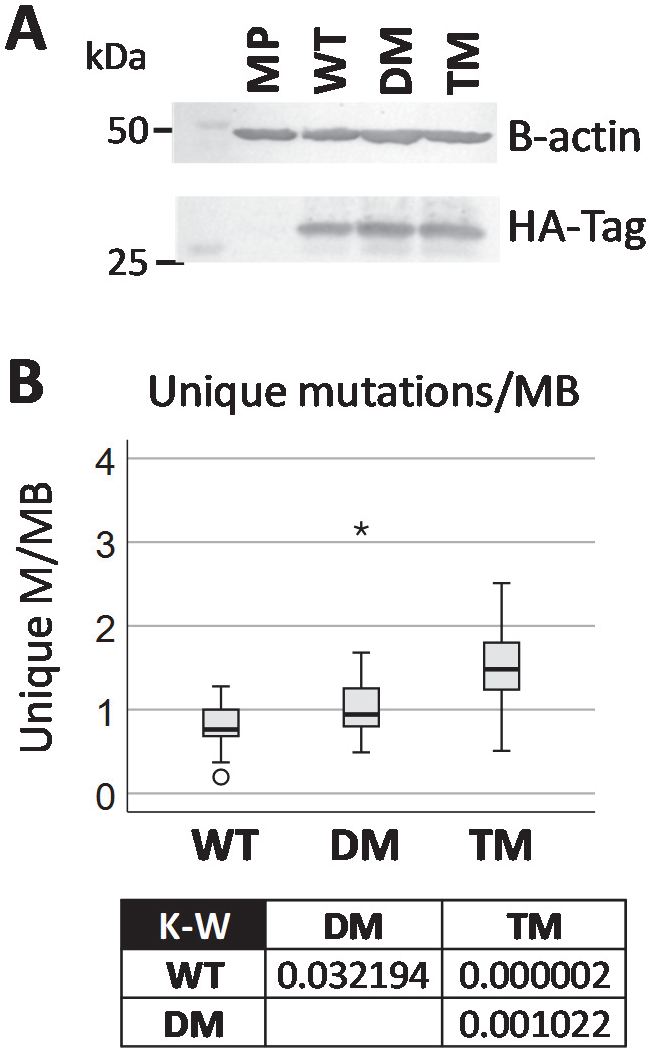
PGRMC1 phosphorylation status effects DNA mutation rate. (A) Western blot of cells following thirty passages showing expression of HA-tag. Beta actin was used as a control. (B) Unique mutations per cell line relative to human reference genome and other genomes in the analysis. Mutations/MB are plotted for 23 chromosomes excluding Y. The box shows *p* values for Kruskal-Wallis (K-W) post-hoc pairwise comparisons.

### PGRMC1 phosphorylation status regulates the NNMT/1-MNA pathway

To further characterize metabolic differences between cells we performed a pilot metabolomics study. In a pilot metabolomics profiling experiment we observed elevated levels of 1-methylnicotinamide (1-MNA) in some but not all experiments (not shown), which we nevertheless chose to explore. 1-MNA is produced by the enzyme nicotinamide-N-methyl transferase (NNMT), which we found to be elevated in TM cells (Figure 6A). NNMT utilizes S-adenosylmethionine (SAM) as a methyl group donor to convert nicotinamide to 1-MNA (Canto et al., 2015) (Figure 6B).

**Figure 6.**
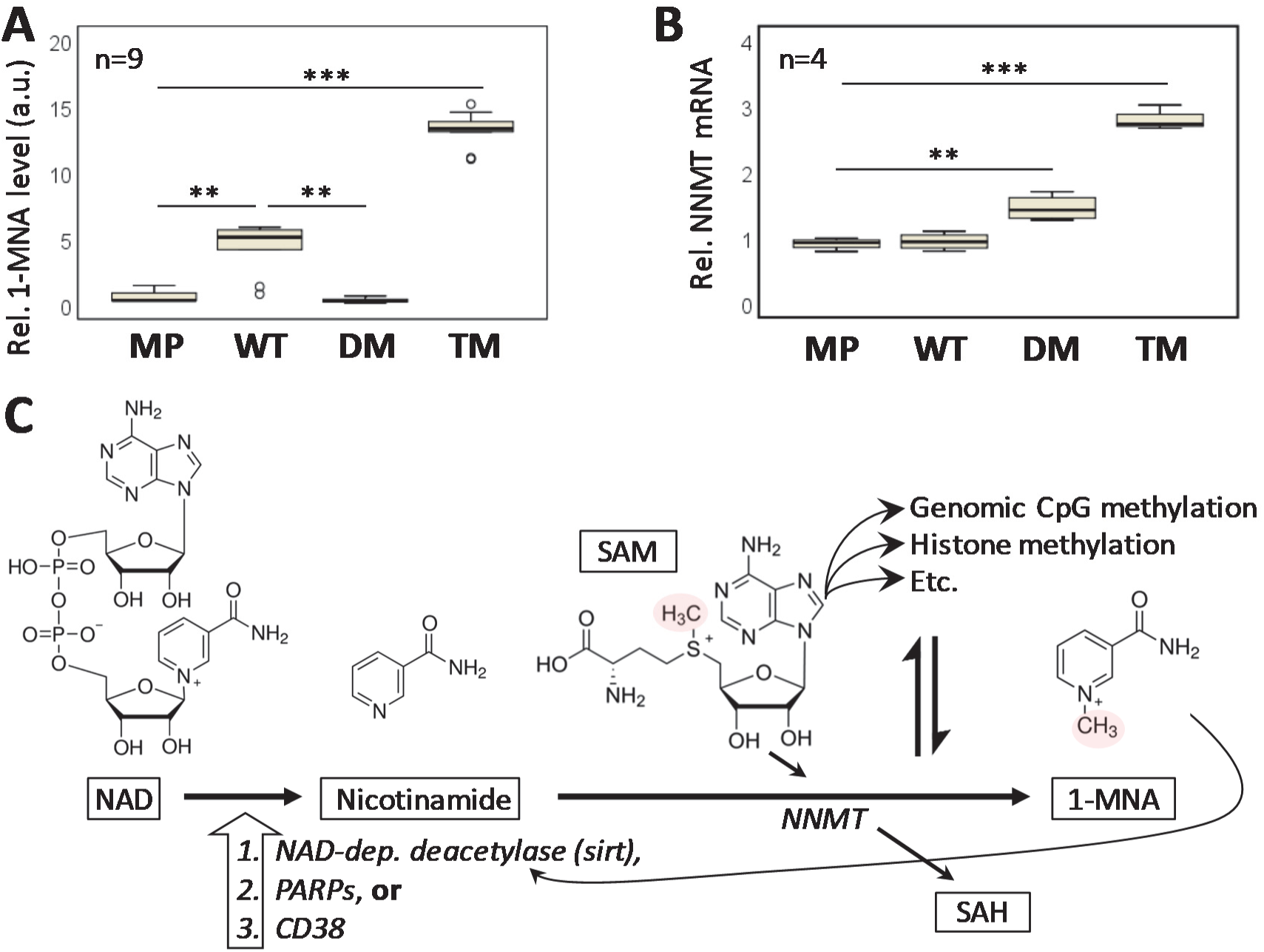
PGRMC1 phosphorylation is associated with NNMT pathway regulation. (A) 1-MNA levels determined by triple quadrupole MS/MS. The y axis represents 1-MNA ion intensity (arbitrary units; a.u.). Samples represent technical triplicates of each of three independently derived stable cell lines or replicates to generate n=9 per condition, or n=3 for MP control. ANOVA post-hoc Tukey’s HSD *p*<0.01 (**) or *p*<0.001 (***). (B) Quantification of NNMT mRNA levels in the four cell types by RT-PCR. Quantification cycle (Cq) results were quantified using the 2^ΔΔCq^ method using beta actin as an internal control. Statistical significance portrayal follows A. (ANOVA post-hoc Tukey’s HSD). (C) Model of the NNMT and 1-methylnicotinamide pathway. Nicotinamide is produced from NAD by NAD-dependent deacetylases (e.g. Sirtuins), PARPS or CD38 (Schultz and Sinclair, 2016). NNMT transfers a methyl group of S-adenosylmethionine (SAM) to nicotinamide to produce 1-methylnicotinamide (1-MNA), producing S-adenosyl homocysteine (SAH). This reaction is in competition with methylation reactions including DNA and histone methylation by SAM (Sperber et al., 2015). 1-MNA can feed back to stabilize sirtuin 1 (Hong et al., 2015).

Liquid chromatography/mass spectrometry (LC/MS) quantification revealed lower levels of SAM in DM and TM cells (Figure S4A). Anti-NNMT shRNA attenuated NNMT levels in TM cells more than two-fold (Figure S4B), which led to expected reduced levels of 1-MNA (Figure S4C). Contrary to the hypothesis that increased NNMT activity would reduce SAM levels as proposed for pluripotent hESCs (Sperber et al., 2015), SAM levels were not significantly altered by NNMT attenuation (Figure S4D). We also treated DM cells with 1-MNA, which caused a morphological conversion from predominantly rounded to predominantly elongated form (Figure S4E), suggesting that NNMT activity may contribute to the morphological differences between elongated MP and WT cells on the one hand and rounded DM and TM cells on the other. Other cell-types showed no significant change in cell morphology upon 1-MNA treatment (not shown).

### PGRMC1 phosphorylation status regulates genomic methylation

Because of the reported effects of NNMT activity on global methylation levels (Sperber et al., 2015), we assayed genomic CpG methylation levels. Hierarchical clustering of samples displayed striking separation of the cell groups, suggestive of highly divergent patterns of methylation among the cell types. All independently derived stable cell lines of each PGRMC1 state clustered closely to each other, and distant from other cell conditions (Figure 7A).

**Figure 7.**
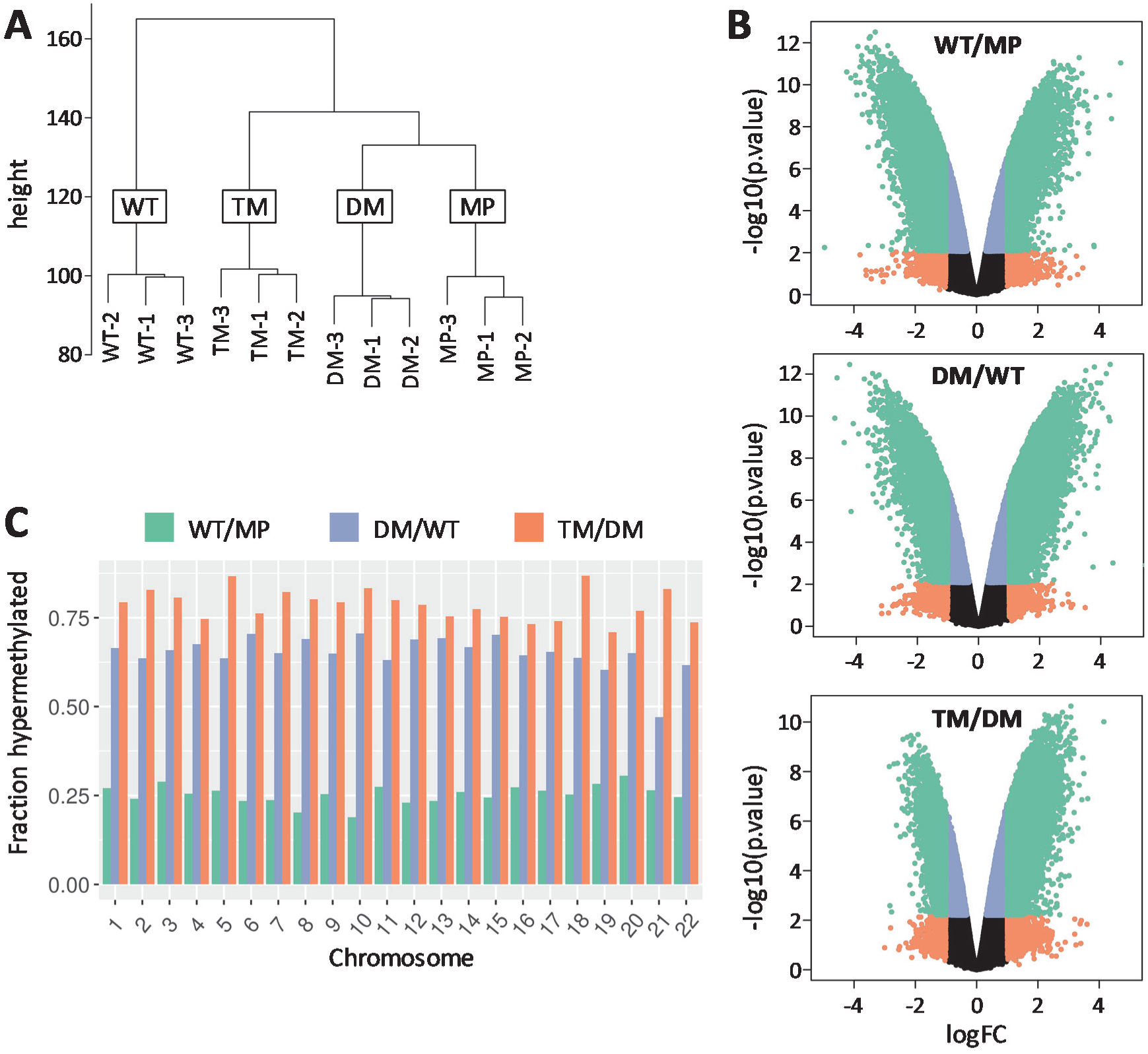
PGRMC1 phosphorylation mutants elicit distinct patterns of genomic methylation. (A) Hierarchical clustering of methylation data from all cell lines sampled in triplicate. Clustering was applied using 520,031 probes with a coefficient of variation > 0.10. Height is calculated using the UPGMA method in the Lumi package and equates to the average distance between two members of two groups. (B) Volcano plots of the WT/MP, DM/WT and TM/DM comparisons. The log2 fold change (FC) (x axis) is plotted relative to the log10 *p*-value (y axis) to visualize the magnitude of the most significant features. Probes with an adjusted *p*-value < 0.05 and absolute log FC > 1 are plotted in green. Probes with an absolute logFC > 1 that were not significant are plotted in orange. Probes with a logFC < 1 that were significant at adjusted *p*-value < 0.05 are plotted in blue. Black probes were not significant. (C) Significant probes in each differential methylation comparison were extracted and the fraction of hypermethylated or hypomethylated probes per chromosome calculated. Methylation levels are higher in TM cells across all chromosomes.

We observed a highly significant number of differentially methylated probes (Figure 7B). In the TM/DM comparison (differ by Y180 state) we observed a shift towards hypermethylation that was not as pronounced in WT/MP (differ by transfection of pcDNA3-1_PGRMC1-HA and hygromycin stable selection) or DM/WT (differ by PGRMC1-HA S57/S181 status). We then considered the percentage of differentially methylated probes which were hypermethylated in each comparison and noticed a clear increase in the progression from WT/MP, DM/WT to TM/DM (Figure 7C). While the WT/DM effect could be due to both PGRMC1-HA expression and/or the stable antibiotic selection process, the differences between DM/WT and TM/DM were due solely to PGRMC1 mutation status. The overall level of CpG methylation was higher in TM cells across all chromosomes (Figure 7C). Together, the results suggest a global increase in net methylation in this system, being highest in TM cells.

Significantly differentially methylated probes were next assessed by genomic context region based on the EPIC manifest designations for relation to CpG island (Island; Open Sea, North Shelf, North Shore, South Shore, and South Shelf) because DNA methylation levels are known to depend upon genomic context (Jones, 2012). The percentage of significantly hypermethylated probes was similar for all PGRMC1-HA-mutation states (WT, DM, TM), which differed from MP (Figure S7A, left). By contrast, there were marked differences in the distribution of hypomethylated probes due to PGRMC1-HA mutation status, with DM and especially TM cells exhibiting elevated levels of hypomethylation at CpG islands and both North and South Shore sites, and reduced levels of hypomethylation (increased methylation) in Open Sea regions (Figure S7A, right). This is consistent with increased levels of Open Sea methylation (in excess to Island and Shore sites) leading to the net increased methylation of TM cells, with a minority of local demethylation sites in the vicinity of CpG Islands at activated genes. See also the volcano plot of Figure 7B, TM/WT comparison, where positive fold change (logFC >0) is in excess.

The increased level of hypomethylation at Islands and Shores in TM cells was not detectable between 5’ untranslated and 3’ untranslated regions of annotated coding genes (Figure S5). A proportion of cell-type and promoter-associated enhancers exhibited reduced hypermethylation or increased hypomethylation in the TM/DM comparison (Figure S6). This reduced level of enhancer methylation, normally associated with gene activation, occurred in the background of bulk overall increased CpG methylation in TM cells. Since TM and DM cells differ only in the presence of Y180 of the exogenous PGRMC1 protein, we conclude that Y180 phosphorylation status appears to specifically influence the genomic methylation status of cell-type specific enhancers.

We performed Kyoto Encyclopedia of Genes and Genomes (KEGG) pathways and Gene Ontology (GO) enrichment analysis of the methylomics results, using probes from Island or Shore genomics regions only. For KEGG we separated the data into separate hyper- and hypo-methylated gene sets. The KEGG results are presented in Figure S7B, and full GO and KEGG results are provided as File S1. Each mutation state induced specific unique suites of detected pathways. Most significantly differential KEGG pathways were detected by all three cell comparisons.

Across all comparisons, the most significant enriched GO terms included some associated with embryology and developmental processes (Table S2, File S1). KEGG results reflected the activities of signal pathways, cell-extracellular interactions, and cancer biology (Table S2, File S1). E.g., for the TM/DM cell comparison, which differs in only the presence of phosphate acceptor oxygen of PGRMC1-HA Y180, KEGG results indicated that the most hypomethylated pathway in TM cells was PI3K-Akt (path:hsa04151). The PI3/Akt pathway was also among the top 5 in the hypermethylated data set for that comparison, and was significantly differential in all three comparisons (File S1). In an accompanying manuscript we predicted reduced PI3K/Akt activity in TM cells on the basis of proteomics pathways enrichment, where the pathway was also significantly differential for all cell comparisons, and we demonstrated reduced phosphorylation of Akt substrates in TM cells (Thejer et al., 2019).

By way of illustrative example, we further explored the biology underyling the pathways analysis by considering the unique KEGG pathways identified for both hyper- and hypo-methylated genes from Figure S7A. 13 hyper-methylated and 7 hypomethylated pathways were unique to the TM/DM comparison (Figure S7B). Their identities are given in Table S3. Hypermethylated pathways included processes related to proteolysis and bacterial infection as well as DNA base excision repair and mismatch repair involved in averting mutations. The latter is striking because we also detected protein abundances involved in pathways associated with DNA damage (p53 effectors, and ATM/ATR), to be reduced in TM cells by proteomics (Figure 4), with accompanying elevated mutation rate in TM cells (Figure 5), which is fully consistent with the methylomics pathways biology. These circumstances suggest strongly that the study reflects true biology related to PGRMC1 Y180 function.

Whereas NNMT was more abundant in TM cells (Figure 6A), CpG sites in its gene were hypermethylated (not shown) as were the sites in many coding genes (Figure S5). Histone methylation status, which we have not assayed, could be influential in determining gene expression levels, or a hypomethylated enhancer may activate transcription despite CpG methylation of the immediate gene locus. Approximately 80% of the genes encoding the 243 differential proteomics heat map of the accompanying paper (Thejer et al., 2019) were significantly differentially methylated between one of the comparisons (not shown), suggesting that epigenetics may contribute greatly to the phenotypes observed for these cell types.

## DISCUSSION

We show that changes in posttranslational modification of PGRMC1 are involved in the maintenance of genomic integrity. Future research should examine whether this involves previously reported PGRMC1 involvement with G0/G1 checkpoint and/or spindle association (Cahill et al., 2016a; Griffin et al., 2014; Juhlen et al., 2018; Luciano and Peluso, 2016; Peluso et al., 2014; Terzaghi et al., 2016).

Perturbation of the regulation of Y180 phosphorylation motif, which is near both S57 and S181 in the folded protein (Cahill, 2007; Cahill et al., 2016b), can induce considerable changes in cellular plasticity. This may be akin to reprogramming aspects of multicellular regulatory mechanisms back to a state more typical of early embryogenesis, which is highly relevant to cancer. It may indeed be associated with the altered prognosis of triple negative breast cancers (Neubauer et al., 2008). However our most important finding is that PGRMC1 Y180 affects pathways associated with animal embryology, seemingly by targeting the genomic methylation status of CpG sites in a subset of cell-type and promoter-associated enhancers.

Vertebrate embryology relies heavily on epigenetic histone and genomic methylation to direct cell plasticity between generations and in response to stimuli to determine the differentiation status of cells (Kiefer, 2007). Y180 (and Y139) appeared in animal evolution in the common ancestor of Cnidaria and Bilateria (Hehenberger et al., 2019), concomitantly with the organizer that drives the embryological tissue differentiation processes. PGRMC1 Y180 was therefore evolutionarily acquired before the appearance of chordate organs, and the embryological differentiation processes that give rise to them, which immediately suggests how PGRMC1 may affect cancer cell plasticity. The increase of hypermethylated CpG sites observed in TM cells (Figure 7, Figure S5) is indeed reminiscent of the characteristic increase in hypermethylation associated with both hESCs and induced pluripotent stem cells (iPSCs). The genome subsequently becomes increasingly hypomethylated during embryogenesis and somatic tissue differentiation (Nishino and Umezawa, 2016). It is as if differentiation processes depended upon the acquisition of Y180 and the organizer by the Cnidaria/Bilateria common ancestor (Hehenberger et al., 2019), and, upon disturbance with Y180 functions in TM cells, the genomic CpG methylation pattern reverts to a phenotype reminiscent of the undifferentiated state of that ancestral evolutionary stage, prior to the appearance of deuterostome and chordate tissues and body plan.

NNMT regulates pluripotency in naïve hESCs, where it was proposed to deplete cellular SAM levels and pleiotropically lead to reduced histone methylation, including some genes in the Wnt pathway whose concomitant elevated expression affected pluripotency (Sperber et al., 2015). PGRMC1 involvement in hESC pluripotency (Kim et al., 2018) suggests that it could be mechanistically involved in the epigenetic modulation that regulates hESC pluripotency (Sperber et al., 2015). Perhaps the most striking result we observed was that each PGRMC1 phosphorylation mutant condition elicited a distinctly unique pattern of genomic methylation (Figure 7), consistent with PGRMC1 playing a central role(s) in regulating genomic methylation, with each phosphorylated isoform capable of dynamically engendering different end-points.

As a target of Wnt signaling, TCF/LEF appeared very early in animal evolution, being absent from the protist Choanoflagellate sister group to metazoans, but present in sponges (Cadigan and Waterman, 2012). By the time the common ancestor of Cnidaria (corals, jellyfish, etc.) and Bilateria (bilaterally symmetrical animals) evolved, embryological gastrulation initiated from the animal pole under control of Wnt/β-catenin signaling, inducing an axial organizer whose vertebrate descendant is known as the Spemann-Mangold organizer. This initiates the orchestrated series of events where specific gene programs are activated by transcription factors that trigger differentiation into specialized cell types (Cadigan and Waterman, 2012; Genikhovich and Technau, 2017; Kraus et al., 2016; Nielsen et al., 2018).

These processes must represent evolutionary novel metazoan functions. The correlation of appearance of the Y139/Y180 combination and organizer function (Hehenberger et al., 2019) is consistent with the involvement of PGRMC1 in this process. This new hypothesized PGRMC1 functionality would seem to be superimposed upon more ancient PGRMC1 functions, such as sterol synthesis and steroid responsiveness. In support of this hypothesis, all of our mutants responded similarly to cyP450-requiring P4-dependent resistance to doxorubicin-induced death, as well as to AG-205 toxicity (Figure 3).

PGRMC1 is required to maintain the pluripotency of embryonic stem cells by influencing the p53 and Wnt/β-catenin pathways (Kim et al., 2018). In an accompanying paper (Thejer et al., 2019) we show that TM cells exhibit lower levels of PI3K/Akt activity, and lowered levels of GSK3-β phosphorylation on an Akt target site. The canonical Wnt pathway is activated when β-catenin translocates to the nucleus to act as a transcription factor after activation by Wnt ligands. Active GSK3-β phosphorylates β-catenin, targeting it for polyubiquitination and proteasomal degradation (Hermida et al., 2017), and Akt phosphorylation of GSK3-β inhibits GSK3-β activity (Hermida et al., 2017).

While our cells are not embryonic, lack of Y180 phosphorylation and low PI3K/Akt activity could correlate with increased GSK3-β activity, and lower β-catenin levels which would attenuate Wnt signaling via TCF/LEF. PGRMC1 has been associated with negative regulation of TCF/LEF in hGL5 granulosa/luteal cells (Peluso et al., 2012) and hESCs (Kim et al., 2018). Further experiments are required to explore this concept further in this and other cell models, which should be prioritized.

The propensity for animals and their cells to age was correlated with the loss of pluripotent stem cells in the adult: an evolutionary necessity to establish larger collections of clonally communal but functionally specialized cells with diverse differentiation states (Petralia et al., 2014). That entire biology was associated with the evolution of the tissue and cell-type heterogeneity of bilateral animals, which started with the evolutionary appearance of the organizer (Petralia et al., 2014). The possible involvement of PGRMC1/PGRMC2 phosphorylation in cell cycle control mechanisms governing entry to G_0_ and senescence (Griffin et al., 2014; Peluso et al., 2014; Shih et al., 2019; Terzaghi et al., 2016) may accordingly be related to an overarching function associated with the reduction of the immortality potential of primitive eukaryotes, and the necessary evolution of a systematics of differentiation status and replicative control in bilateral animals (Petralia et al., 2014). This hypothesis merits future examination.

The NNMT pathway has been linked to lifespan and aging (Calvani et al., 2014; Neelakantan et al., 2019; Schmeisser et al., 2013; Schmeisser and Parker, 2018), to sirtuin protein stabilization in positive a feedback loop to the pathway in Figure 6B involved in energy regulation and obesity (Hong et al., 2015; Kraus et al., 2014; Strom et al., 2018), and to epigenetics in cancer and stem cell status (Jung et al., 2017; Sperber et al., 2015; Ulanovskaya et al., 2013; Xie et al., 2014) (reviewed by (Pissios, 2017)). We show that PGRMC1 Y180 modulates this system.

In mining our proteomics data for enzymes related to the NAD/NNMT/1-MNA pathway (Figure 6B), we noted that nicotinamide phosphoribosyltransferase (NAMPT) was one of the relatively few proteins less abundant in TM compared to both DM and WT (Figure S3). NAMPT catalyzes a competing reaction to NNMT, whereby nicotinamide is converted to nicotinamide mononucleotide, which is then transformed into NAD^+^ by nicotinamide/nicotinic acid mononucleotide adenyltransferase. NAMPT is the rate-limiting enzyme in this pathway, and regulates sirtuin activity by modulating NAD^+^ levels (Revollo et al., 2004). This entire pathway is related to aging processes (Dai et al., 2018), which were acquired by bilaterian animals along with tissue differentiation and cell cycle control mechanisms to evade cancer (Petralia et al., 2014). Indeed, these are the very types of biology with which PGRMC1 function is here implicated: suggesting that PGRMC1 phosphorylation could be involved in the mechanism(s) of animal aging. Considering the long co-evolutionary history of Bilateria and PGRMC1 tyrosine phosphorylation sites, this merits future study.

Our results imply that the ability to interchange between phosphorylated states at T178, Y180, S181 and S190 (Cahill et al., 2016b) will be critical in supplying correct PGRMC1 function at appropriate times for the cell cycle and state of cell differentiation during early embryology. Clearly, Y180 is important for PI3K/Akt activity (Thejer et al., 2019), a major pathway associated with aging (Wrigley et al., 2017).

S181 has been discussed in the literature as a consensus CK2 site (Cahill et al., 2016a). It must be noted in this context that PGRMC1 phosphorylation at S181 was unaffected by knockout of CK2 kinase activity in C2C12 mouse myoblast cells (Franchin et al., 2018), indicating that at least in those cells CK2 was not responsible for S181 phosphorylation. Willibald et al. (Willibald et al., 2017a) recently used an antibody against PGRMC1 phospho-S181 (pS181) to argue that S181 phosphorylation is essential in PGRMC1 activation, as well as discussing the presence of phosphorylated PGRMC1 in immunohistochemistry of tissue biopsies (Willibald et al., 2017b). A caveat to these results is that the peptide epitope used to generate the antibody (which was designed and commissioned by M.A.C. while at ProteoSys AG, (Neubauer et al., 2009; Neubauer et al., 2008)), corresponded to a CK2 consensus site. CK2 is a promiscuous kinase which gives rise to 10% - 20% of the human phosphoproteome (Pagano et al., 2006). On Western blots of breast cancer whole cell proteins, that antibody generated a low background smear of signals across many molecular weights, in addition to recognizing exogenously expressed PGRMC1 but not a S181A mutant (not shown). More rigorous characterization of the anti-pS181 antibody specificity would be required to exclude that the immunohistochemistry of tissue biopsies was not detecting increased levels of non-PGRMC1 CK2 phosphorylation sites.

## Conclusions

This work suggests that PGRMC1 has been a key yet hitherto unrecognized protein involved in the early evolution of animals, and that its phosphorylation status is capable of influencing biological processes which can destabilize the ordered harmony of the metazoan multicellular collection of functionally specialized cells, with imperatively relevant connotations for human disease. Our work was inspired by the discovery that PGRMC1 phosphorylation differs between breast cancer subtypes differing in estrogen receptor expression, that also exhibit starkly contrasting levels of patient survival (Neubauer et al., 2008). It is wholly feasible that characterization of the signal network surrounding PGRMC1 could lead to novel treatments for cancer, and other diseases involving PGRMC1 (Cahill et al., 2016a). That aspiration could have been greatly advantaged (there has been more than a decade hiatus in experimental results publication due to lack of resources) if the Australian national competitive grants system had supported this obviously innovative, highly significant, and demonstrably valid (Cahill et al., 2007 (Priority 20050926); Cahill, 2007; Neubauer et al., 2008) work after M.A.C.’s relocation to Australia in 2008.

## ACKNOWLEDGEMENTS

This work has received no direct Australian competitive grant support since M.A.C.’s relocation to the country in 2008. The present results have been compiled largely due to the generosity of collaborating authors, and by the PhD stipends of B.M.T. and P.P.A. Research was primarily supported by Charles Sturt University (CSU) internal funds to M.A.C., J.A.J. and L.A.W. B.M.T. was supported by a PhD scholarship from the Ministry of Higher Education and Research, through the University of Wasit, Iraq. E.M.G. acknowledges partial support of Australian Research Council award CE140100003. E.J.N. acknowledges the support of the Australian Research Council (DE120102687) and the Ramaciotti Foundation (ES2012/0051).

## AUTHOR CONTRIBUTIONS

Conceptualization, M.A.C.; Methodology, M.A.C., P.P.A, B.M.T., E.M.G., M.G., E.J.N, L.A.W., P.A.W, and M.P; Formal Analysis, B.T.M., P.P.A., S.D.P., R.H.T., P.A.W., M.L., M.G., L.G., T.F., T.F., M.P. and M.A.C.; Investigation, B.M.T., P.P.A., S.L.T., J.F., P.A.W., L.A.W., S.G. and M.W.; Resources, B.M.T., M.P., E.J.N., H.N., T.F., E.M.G, and L.A.W.; Writing – Original Draft, M.A.C., B.M.T.; Writing – Review & Editing, M.A.C., B.M.T., P.P.A., M.P., E.M.G; Supervision, M.A.C, J.A.J., and E.J.N.; Project Administration, M.A.C.; Funding Acquisition, M.A.C., L.A.W., J.C.Q., E.M.G., H.N., T.F., and E.J.N.

## STAR METHODS

### Establishment of cell lines and cell culture

Construction of cell lines and culture conditions is described in the accompanying publication (Thejer et al., 2019).

### NpFR1 measurement of cytoplasmic redox status

Intracellular redox state measurement by fluorescent redox sensor NpFR1 was performed by flow cytometry as described for NpFR2 (Thejer et al., 2019). We have repeated the result of Figure 2 on numerous occasions. The actual ratio of low to high abundance distribution for a cell type tends to vary between experiments, however the relative proportions of cells in the low fluorescent population between cell types remains relatively constant (MP > DM > WT > TM).

### Genomic DNA sequencing

Genomic DNA for sequencing was isolated from one subline of each cell condition. Passaging was repeated every 4 days for thirty passages, with media changes every 48 hours. To extract genomic DNA after thirty passages, approximately 5×10^6^ cells were centrifuged for 5 min at 300 x g. The cells were suspended in 200µL phosphate buffer saline. The purification of total DNA from cells was performed using the DNeasy Blood & Tissue Kit (Qiagen, C.N 69504) following the manufacturer’s protocol. The concentration of the dsDNA was measured using a Qubit dsDNA Hs Assay Kit (Life Technologies, Ref Q32851). The samples were sent to Kinghorn Centre for Clinical Genomics (Garvan Institute of Medical Research, 384 Victoria St, Darlinghurst, NSW 2010 Australia) for whole genomic sequencing. DNA for methylation was extracted the same way except we used passage 10 cell lines, consisting of triplicate cultures of MP cells, and respective lines 1-3 of each PGRMC1-HA mutant condition (WT1, WT2, WT3, DM1, DM2, DM3, TM1, TM2, TM3).

### MTT cell viability assays for effects of P4 on the dox and AG-205 responses

To assay for proliferative and protective effects of P4, 10^4^ cells were seeded per 96 well plate well. For each cell type, n=6 replicates were allowed to adhere for 3 hours after which viable adherent cells were quantified by 3-(4,5-dimethylthiazolyl-2)-2,5-diphenyltetrazolium bromide (MTT) assay. Previous experiments had shown adherence to be complete after 1.5hr (data not shown). Identical replicates were incubated overnight in complete DMEM. After 23 hours the media was exchanged for complete DMEM containing either 1µM P4 or DMSO vehicle control. After a further 1 hour incubation (t0) the media was exchanged for media containing either 1µM P4 or DMSO vehicle control (to assay for proliferative effects, Figure 2A), or with varying concentrations of doxorubicin (dox) (Figure 2B-C) or AG-205, +/- P4 (Friel et al., 2015). Cells were incubated for a further 24hr followed by viable cell quantification by MTT assay. The media was discarded, cells were washed with warm PBS and incubated with 100μL of 0.5 mg/mL MTT (Sigma-Aldrich M2128) in phenol red free media (Sigma-Aldrich D1145). After 3 hours incubation at 37°C and 5% CO_2_, the media was removed and 100μL of DMSO (EMD Millipore, 317275) was added to each well to solubilize the Formazan crystals. The cells were incubated for 1 hour followed by mixing, and absorbance was read at 570nm using a plate reader (Bio-Strategy P/L, Campbellfield, Vic., Australia). The percentage of viable cells was estimated by normalizing the absorbance of the treated or untreated cells to the average values from panel A.

### NNMT shRNA attenuation

We established an anti-NNMT sh-RNA-expressing TM cell line by lentiviral transduction and puromcyin stable selection. Anti NNMT shRNA lentiviral production was as described using mission TRC2-pLKO-Puro series Lentiplasmid (SHCLND, Sigma-Aldrich) scramble shRNA (scr-sh) (Thejer et al., 2019) and shRNA targeting NNMT (Sigma-Aldrich TRCN0000294436, TGCAGAAAGCCAGATTCTTAA). WT and TM cell lines were seeded in 24 well plates until they reached approximately 50% confluency followed by transduction with shRNA virus particles and selection as described (Thejer et al., 2019).

### RT-PCR

RNA was extracted by total RNA mini kit (Bio-Rad, 7326820) according to the manufacturer’s instructions. cDNA was synthesized in C1000 Touch™ Thermal Cycler using cDNA synthesis kit (Bio-Rad, 1708890) following Bio-Rad’s recommended protocol. Real-time qPCR was performed on a CFX96 Touch™ Real-Time PCR Detection System utilizing iTaq™ Universal SYBR^®^ Green Supermix (Bio-Rad, 1725120). NNMT Primers for qPCR were synthesized at Monash University and the sequences were obtained from (Parsons et al., 2011). NNMT primers used in this experiment were Forward sequence 5’-GAATCAGGCTTCACCTCCAA-3’ and Reverse sequence 5’-CCCAGGAGATTATGAAACACC-3’. Actin was used as an internal control Forward sequence 5’GACGACATGGAGAAAATCTG-3’ and Reverse sequence 5’ATGATCTGGGTCATCTTCTC-3’.

### Western Blot

Cells were lysed with radioimmunoprecipitation assay buffer (RIPA buffer) (Sigma Aldrich, R0278) supplemented with protease and phosphatase inhibitor cocktail (Thermoscientific, 88668). After scraping the cells, they were centrifuged at 8000 rpm for 20 minutes (Hermle Centrifuge Z233 M-2) at 4^°^C. 20µg protein of thirty passage number of MP, WT, DM, and TM were loaded into the wells of a 12.5% SDS-PAGE gel and set to run at 150V for 45 min on Power Pac. Transferring the protein from the gel to a PVDF membrane occurred under wet transfer with 1x Towbin buffer at 40V for 3 hours in an ice bath. The membranes were incubated with 1:2000 diluted Beta-Actin (Sigma Aldrich, AS441) and 1:2000 HA-Tag (Sigma Aldrich, H3663) primary antibodies in blocking buffer overnight at 4°C with shaking. Next day, the membranes were washed twice with 0.05% PBS-T and incubated with anti-mouse secondary antibody; horse radish peroxidase (HRP), for 1 hour at room temperature. Next, the membranes were washed twice with 0.05% PBS-T and once with PBS. Colorimetric detection of the bands occurred using tetramethylbenzidine as described (Thejer et al., 2019).

### Reverse phase protein array

Methods for reverse phase protein arrays are described in the accompanying paper (Thejer et al., 2019). The following primary antibodies (provider and reagent number, dilution) were used here: Erk1/2-P-Thr202/Tyr204 (CST 4370, 1:100), MEK1/2-P-Ser217/Ser221 (CST 9154, 1:100), MKK4(SEK1)-P-Ser257/Thr261 (CST 9156, 1:100), Rb (CST 9309, 1:200), Rb-P-Ser807/Ser811 (CST 8516, 1:100).

### Hyperspectral fluorescence microscopy

UV and visible continuous wave epifluorescence microscopy with multiple excitation wavelength ranges from 335 nm to 532nm and measuring emission in three emission wavelength ranges 450+/- 30 nm, 587+/- 17.5 nm and 700 nm long pass were used. Excitation wavelengths are supplied by low cost multi-LED light source optical fibre coupled to the Olympus IX71 microscope. An Andor/Oxford Instruments iXon 885 Electron Multiplying Charged Coupled Device (EMCCD) was used to capture images typically using an EM gain sufficient to lift the low light auto-fluorescence signal above the 17 electrons per pixel per second readout noise but with minimal contribution from clock induced charge. Gain linearity is ensured by using the Real Gain^TM^ technology (Andor/Oxford Instruments). The camera is operated below −70°C such that thermal noise is negligible. Quantum efficiency of the sensor varies from 50-65% across the range of interest from 450-670nm and the intensity is digitized into ∼16K values. Background signal is subtracted from all images which is kept minimal through the use of low fluorescence petri dishes (CELLview, Greiner Bio-One). A single wavelength image may take between 1-5 seconds depending on the sample and wavelength but 1-2 minutes is typically required for the entire stack of images for all spectral channels, where the features for Figure 1A were as follows.

Feature 1: ’Ratio of Channel 4 to Channel 10’,

Feature 2: ’Ratio of Channel 9 to Channel 16’,

Feature 3: ’Ratio Channel 2 to Channel 16’,

Feature 4: ’Ratio of Channel 5 to Channel 14’,

Feature 5: ’Ratio of Channel 2 to Channel 7’,

Feature 6: ’Ratio of Channel 9 to Channel 18’,

Feature 7: ’Ratio of Channel 7 to Channel 15’,

Feature 8: ’Ratio of Channel 7 to Channel 9’,

A description of the channels used and their spectral ranges is given in Table S1. During growth under standard mid-log phase growth conditions, hyperspectral imaging was performed on ∼300 live cells per cell condition (MP, WT, DM, TM) and the mean cellular intensity in each channel was calculated. Pixel correlations between spectral channels were removed using principal component analysis (PCA), followed by a discriminatory projection to maximally separate the four cell groups (Figure 1A). Technical details of this approach are described by Gosnell et al. (Gosnell et al., 2016a). In order to statistically analyze the cluster separations of Figure 1A, an additional LDA projection of cell data was carried out, this time with two classes of cells chosen at a time. Their average spectra were projected onto a common direction identified by this additional LDA. The cell distributions were then tested by the Kolmogorov-Smirnov test.

### Metabolite preparation

Scr-WT, scr-TM, shRNA NNMT-WT, and shRNA NNMT-TM cells were seeded in Greiner Bio-One Tissue Culture Petri Dish 100mm x 20mm (Interpath, 664160). The cells were rinsed with pre-warmed deionised water with brief shaking. The plates were placed on liquid nitrogen for 15 seconds. Cells were extracted with 1 mL of ice-cold extraction solvent (2:2:1methanol-ethanol-water) and harvested with scraper. The contents were transferred to 1.5 mL Eppendorf tube. The tubes were centrifuged at 4°C for 5 minutes at 16,100 x g and supernatant were transferred to a new 1.5 mL Eppendorf tube. The supernatant was filtered and evaporated dry under a gentle stream of nitrogen gas. The residual precipitates were resuspended in 500 µL of 60% acetonitrile/water (ACN/H_2_O) containing m-tyrosine (10 µg/mL) (Sigma, Castle Hill, NSW) as an internal standard.

### Metabolite quantification

Metabolomic analysis was performed using an Agilent 1290 Infinity HPLC system equipped with a quaternary pump, degasser, temperature-controlled column and sample compartment coupled to an Agilent 6470 triple quadrupole (QQQ) mass spectrometer with a Double Jet Stream electrospray ionization (ESI) source (Agilent Technologies, Australia).

The column temperature was maintained at 35°C and the autosampler temperature was set at 4 °C. The electrospray (ESI) source settings were as follows: nebulizer gas, 45 psig; drying gas flow rate, 8.0 L/min; drying gas temperature, 300°C; sheath gas temperature, 350°C, sheath gas flow, 10 L/min; capillary voltage, 4000 V; and nozzle voltage, 0 V. The data were acquired positive ionization mode.

Methylnicotinamide samples and samples from Figure 6A were analyzed with a Kinetex HILIC column (50 mm × 2.1 mm, 2.6 μm particle size, pore size of 100 Å, Phenomenex, CA, USA). The mobile phases were 100% HPLC-grade acetonitrile (A) and 10 mM ammonium acetate in HPLC-grade water adjusted to pH 3 with formic acid (B) at a flow rate of 200 μL/min. The solvent gradient began with 95% ACN for 0.5 minutes, decreasing linearly to 35% ACN over 12 minutes. The gradient was maintained at 35 % ACN for 0.5 minutes and increased to 95 % ACN over 0.1 minutes. The mobile phase composition was then held at 95% CAN for 10 minutes to re-equilibrate the column before the next injection. The total time for the gradient program was 24 minutes.

1-MNA was tentatively identified with *m/z* of 137.0633 Da, and an MS/MS score of 99.74. Samples of authentic 1-methylnicotinamide chloride (1-MNA) (CAS Number 1005-24-9, Sigma-Aldrich # M4627), 2-methylnicotinamide (2-MNA) (CAS Number 58539-65-4, Ark Pharm Arlington Heights, IL 60004, USA, #AK-39636) and N-methylnicotinamide (N-MNA) (CAS Number 114-33-0, Sigma-Aldrich #M4502) were injected at three concentrations (1, 10 and 100 µg /mL) to confirm identification of 1-MNA. The compound was unambiguously identified as 1-MNA with retention time (RT) of 7.33 min and *m/z* of 137.0717, and not 2-MNA (RT 0.96 min, *m/z* 137.0719) or N-MNA (RT 0.97 min, *m/z* 137.0709). The response of the triple quadrupole mass spectrometer to 1-MNA was linear over the range of 10 ng/mL to 1 µg/mL, and the identity was confirmed by MS/MS and HPLC retention time using 2-methylnicotinamide and N-methylnicotinamide standards (not shown).

Cellular extracts except Figure 6A were analyzed with a HILIC-Z column (50 mm × 2.1 mm, 2.6 µm particle size, 100 Å pore size, Agilent Technologies, CA, USA) equipped with a guard column of the same stationary phase. The mobile phases were HPLC-grade acetonitrile:water (95:5) (A) and water (B), both containing 20 mM ammonium formate and 0.1% formic acid (pH ∼3) at a flow rate of 400 µL/min. The solvent gradient started at 100% A and was held for 0.5 minutes, followed by a linear gradient from 100% A to 60% A over 9 minutes. The gradient was maintained at 60% A for 0.5 minutes and returned to 100% A over 0.1 minute. The solvent was then held at 100% A for 5.4 minutes to re-equilibrate the column before the next injection. Injection volume was 2 µL. The total time for the gradient program was 15 minutes.

Analysis of the set of samples was preceded and followed by injection of five concentrations of 1-MNA (10 µg/mL to 1 ng/mL in log steps) and six concentrations of S-adenosyl methionine (SAM) (100 µg/mL to 1 ng/mL) (Sigma-Aldrich, Castle Hill, NSW). In addition, after every 15 samples a blank injection and a mixture of 1-MNA, m-tyrosine and SAM was run to verify that there was no carryover between samples and that retention time and detector sensitivity had not changed during the course of running the samples.

The selective detection and quantitation of 1-MNA was achieved by monitoring the transition of *m/z* 137 → 108 using a fragmentor voltage of 120 V and collision energy of 20 V, with the further requirement that retention time matched that of the authentic compound (ca. 2.7 min). m-tyrosine was quantitated by monitoring the transition *m/z* 182 –> 136 (fragmentor voltage 135 V, collision energy 20 V, RT = 4.9 min) and SAM was quantitated by monitoring the transition *m/z* 399 -> 136 (fragmentor voltage 100 V, collision energy 20 V, RT = 9.1 min). The fragmentor voltage and collision energy values were determined by optimizing system response to authentic 1-MNA, m-tyrosine and SAM. Limits of quantitation were determined by regressing the log(detector response) to log(concentration) response over the range of serial dilutions for each standard, and by observing where the dose/response curve departed from linearity.

### Effect of 1-MNA on morphology

MiaPaCa-2 cells expressing DM PGRMC1-HA were seeded at 25% confluency and cultured overnight. The media was removed and the cells were incubated in the presence of the indicated amount of 1-MNA (Sigma-Aldrich, Castle Hill, NSW) for 24 hours. The 1-MNA stock was made in sterile H_2_O. Five random images were taken per treated culture and assigned random numbers 1-20. These were given to a blinded scorer who decided for each image whether given cells were round or elongated.

### Methylome Assay

Genomic DNA from each cell subline were processed using the Illumina Infinium HD Methylation Assay (EPIC array) which interrogates >850,000 CpG sites. The four cell lines were processed in triplicate equating to 12 samples, being three replicates of MP cells, and independent cell lines 1-3 for each of the PGRMC1 states (Thejer et al., 2019). The array was prepared at the Australian Genome Research Facility (AGRF) following the manufacturer’s instructions. Quality checking of the samples was performed by Nanodrop Spectrophotometer and resolution on a 0.8% agarose gel at 130 V for 60 minutes. 500ng total DNA was bisulfite converted with Zymo EZ DNA Methylation kit (Zymo Research, Irvine, CA, USA), following the manufacturer’s standard protocol. Amplified DNA samples were fragmented and resuspended DNA was loaded onto a BeadChip. The BeadChip was incubated overnight while DNA fragments anneal. After hybridization the array was imaged on the Illumina iScan system.

Methylation data were processed using in R version 3.5.1 (www.r-project.org). Data were assessed using the hg19 build of the human referene genome annotated in the IlluminaHumanMethylationEPICanno.ilm10b4.hg19 bioconductor annotation. Quality control of the array was assessed with the R package Lumi (Huber et al., 2015). Probes on the sex chromosomes, cross-reactive probes and probes with known SNPs at the CpG site were excluded. Following filtering 812,817 probes remained. The cleaned data were normalized with the SWAN normalization, and sample relationships examined. Genome-wide differential methylation analysis was undertaken using Limma (Ritchie et al., 2015). Multiple testing correction was applied using the Benjamini and Hochberg method. Differential methylation in each group was observed on volcano plots and heatmaps.

Due to the excessive quantity of differentially methylated probes with small effects we refined the data used for functional analysis to the top 2000 most variable probes prior to pathways analysis. These probes explained >90% of the total variance among the first two principle components while preserving a reasonable number of intragenic probes. Pathways enrichment was undertaken in R using the *enrichPathway* function in the ReactomePA package (Yu and He, 2016), which implements a one-tailed hypergeometric test for overrepresentation. Multiple testing adjustment was applied with the Bonferroni & Hochberg FDR. KEGG pathways analysis was undertaken using the *kegga* function in the R package in LIMMA. Enrichment was calculated for genes from the top 2000 most variable probes with a *p*-value <0.05. The gene sets were split to those with hypomethylated probes (logFC<0) and those with hypermethylated probes (logFC>0) for enrichment. Enrichment of Gene Ontology (GO) terms was undertaken using the GOstats package (Falcon and Gentleman, 2007). Ontology assignments are calculated using a hypergeometric test for overrepresentation on the significantly differentially methylated probes (*p*<0.05) among the 2000 most variable. Multiple testing adjustments were applied using the FDR. Intersects of common pathways from KEGG and Reactome analyses were assessed using the UpSetR package (Conway et al., 2017).

Methylomics data have been deposited in the ArrayExpress database at EMBL-EBI (www.ebi.ac.uk/arrayexpress) under accession number E-MTAB-8218. Genomic sequencing results are available as NCBI Bioproject Accession: PRJNA400337, SRA Run Selector SRP118430 (https://www.ncbi.nlm.nih.gov/bioproject/PRJNA400337). Further data are available in a supplemental information file available from the journal web page.

## LIST OF ABBREVIATIONS

1-MNA: 1-methylnicotinamide
adjP: Benjamini-Hochberg adjusted p-value
AG-205: AG-205 PGRMC1 inhibitor
CK2: casein kinase 2
cyP450: cytochrome P450
CYP51A: lanosterol 14-alpha demethylase
cytb5: cytochrome b5
Dap1: damage-associated protein 1
DM: PGRMC1-HA S57A/S181A double mutant
Dox: doxorubicin
GLP-1: glucagon-like peptide-1
GO: Gene Ontology
GSK3-β: Glycogen synthase kinase 3-β
HA: hemagglutinin
hESCs: human embryonic stem cells
KEGG: Kyoto Encyclopedia of Genes and Genomes
LC/MS: Liquid chromatography/mass spectrometry
LDL: low density lipoprotein
LDLR: low density lipoprotein receptor
MAPR: membrane-associated progesterone receptor
MP: MIA PaCa-2 cells
mPRα: membrane progestin receptor α
NAMPT: nicotinamide phosphoribosyltransferase
NNMT: nicotinamide-N-methyltransferase
NpFR1: Naphthalimide-flavin redox sensor 1
NpFR2: Naphthalimide-flavin redox sensor 2
P4: Progesterone
PGRMC1: Progesterone Receptor Membrane Components 1
PGRMC1-HA: hemagglutinin-tagged PGRMC1
RPPA: reverse phase protein array
SAM: S-adenosylmethionine
SH2: Src homology 2
SH3: Src homology 3
TCF/LEF: T-cell factor/lymphoid enhancer factor
TM: PGRMC1-HA S57A/Y180F/S181A triple mutant
WT: PGRMC1-HA wild-type

## Supplementary Figures

**Figure S1.**
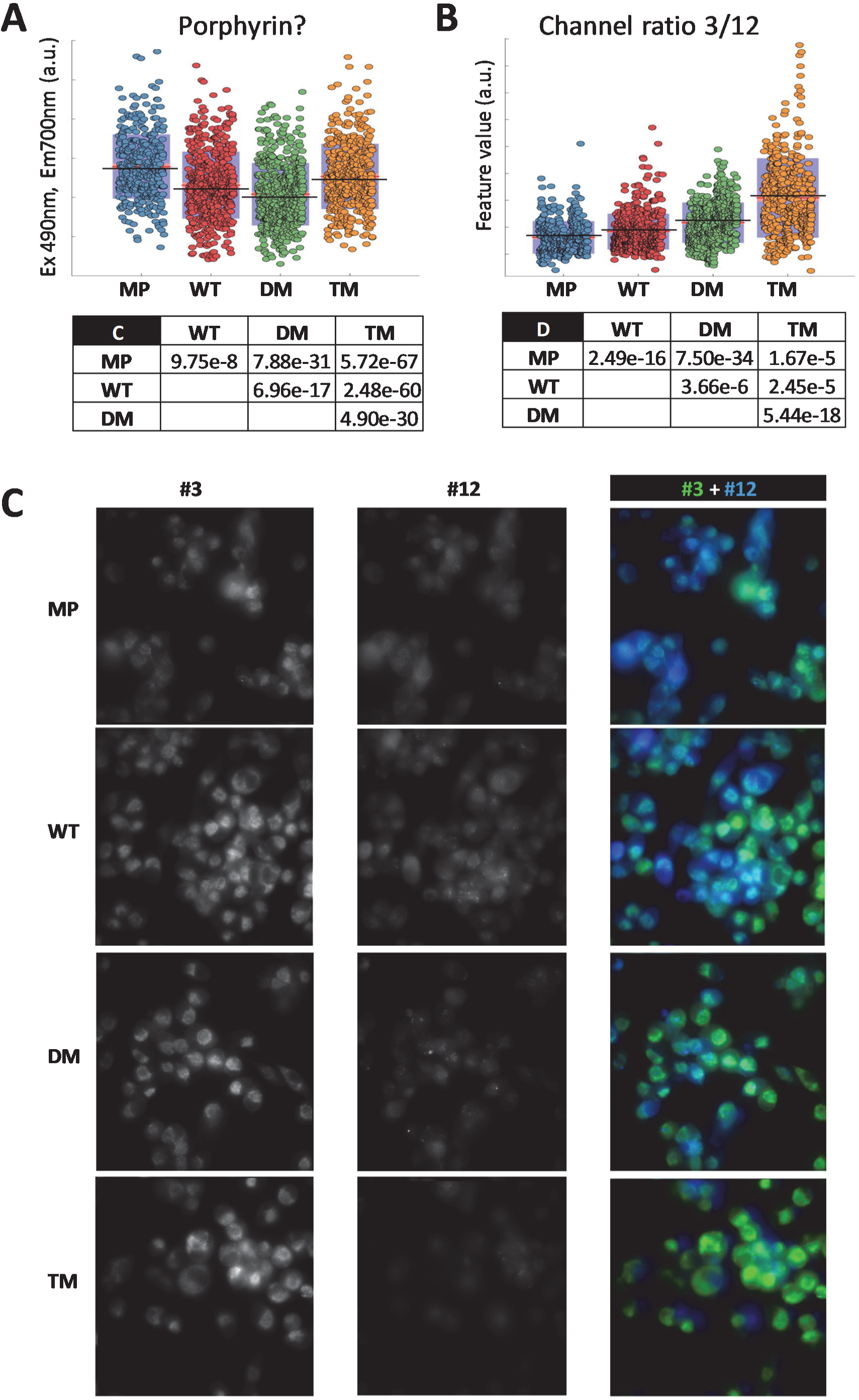
Representative hyperspectral autofluorescence cell images from the measurements. Related to Figure 1. (A) Mean cellular intensity of hyperspectral autofluorescence channel 18 [495nm(Ex), 700nm(Em)], which may reflect porphyrin or protein-bound red-shifted flavin emission (Schneckenburger et al., 1989), is significantly affected by PGRMC1-HA phosphorylation status. The table provides Kolmogorov-Smirnov test p values from pair wise comparisons. (B) The ratio of hyperspectral autofluorescence channels 3 [375nm(Ex), 450nm(Em)] to channel 12 [435nm(Ex), 587nm(Em)] differs significantly between cells. The table follows C. (C) Individual channels #3 and #4 from (B) as listed in Table S1 (left) and the same two channels superimposed (right).

**Figure S2.**
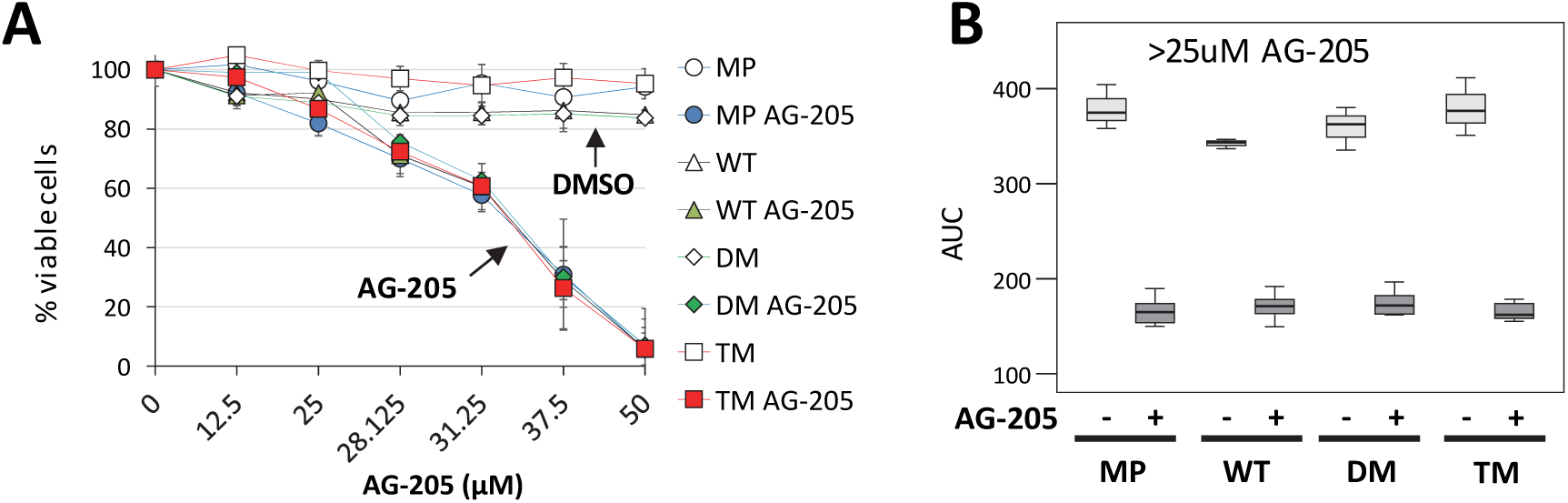
PGRMC1 phosphorylation mutants do not affect AG-205-induced death. Related to Figure 3. (A) AG-205-induced cell death is unaffected by PGRMC1 phosphorylation status. Cells were incubated in the presence of the indicated AG-205 concentrations (n=8: 3x line 1, 3x line 2, 2 x line 3) or DMSO vehicle control (n=3: 1x each cell line), and percentage viable cells was calculated relative to untreated cell controls (n=9: 3 replicates per cell line) using MTT assay. (B) AUC results for values from A greater than 25μM reveal no significant differences in response to AG-205 treatment between cell lines (p>0.85, post-hoc Bonferroni after 1 way ANOVA for AG-205 treatment). The apparently greater survival of DM cells at 25 μM AG-205 observed in this panel was never observed in multiple other repeat experiments.

**Figure S3.**
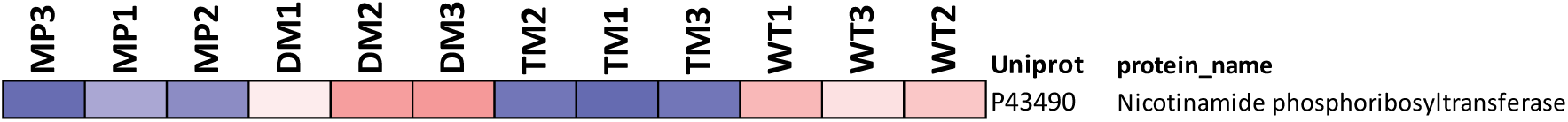
SWATH-MS proteomic quantification of Nicotinamide phosphoribosyltransferase. Related to Figure 6. The figure shows the abundance profile of P43490 nicotinamide phosphoribosyltransferase (NAMPT) from the SWATH-MS proteomics quantification of the accompanying manuscript (Thejer et al., 2019).

**Figure S4.**
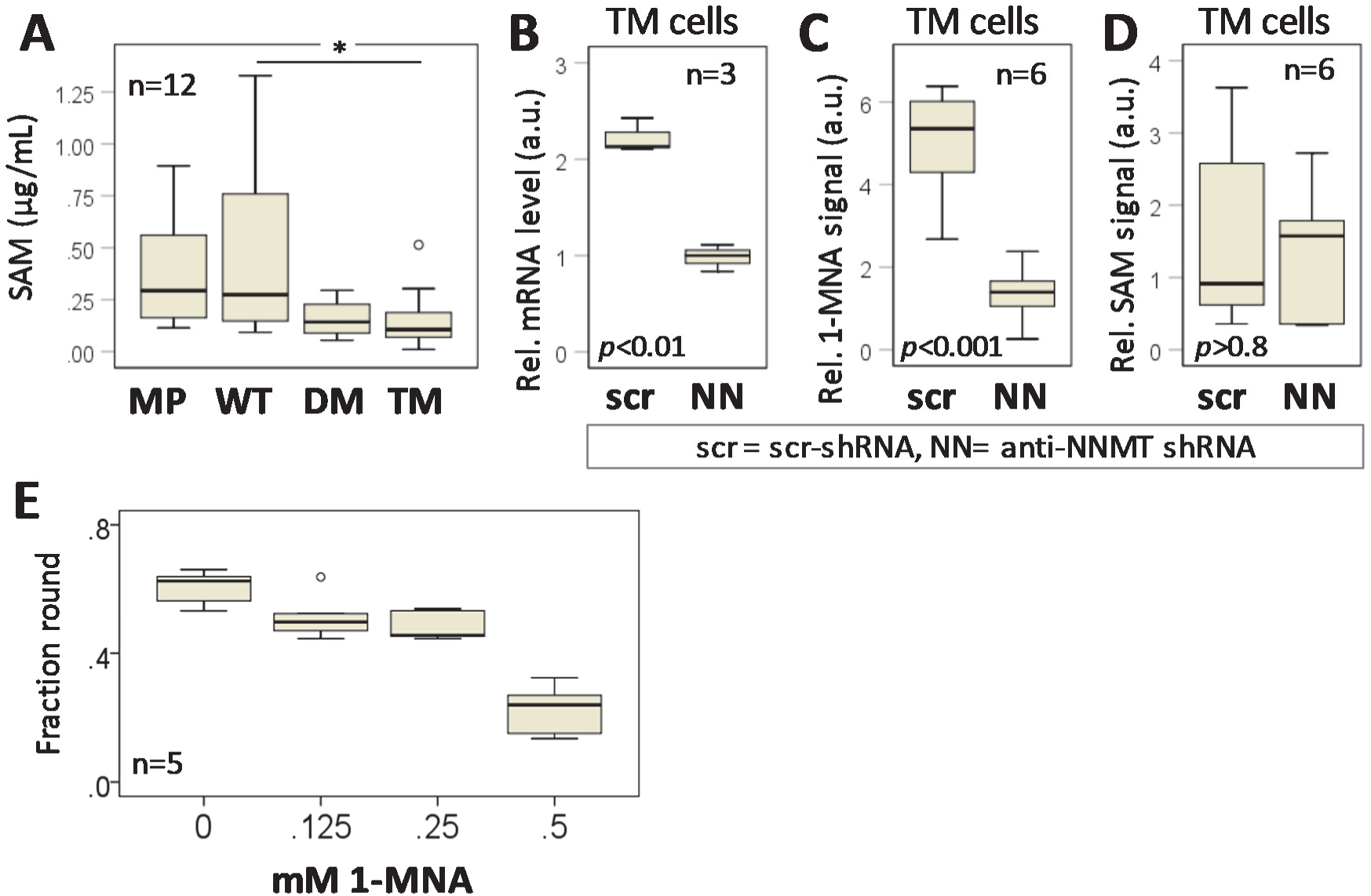
SWATH-MS proteomic quantification of Nicotinamide phosphoribosyltransferase. Related to Figure 6. (A) Metabolomics quantification of S-adenosyl-Methionine (SAM) levels for the indicated cell lines, representing n=12 (four technical replicates for each of three independent cell lines per PGRMC1 condition). (B) RT-PCR quantification of NNMT mRNA levels after treatment by an NNMT-specific shRNA (NN) or a random scramble shRNA control (scr) in TM cells stably expressing lentiplasmid-driven shRNAs. RT-PCR Methods follow A. p <0.003 (2-tailed T test). Methods follow Figure 6B. (C) 1-MNA ion intensities in cells from B, determined following methods from Figure 6A, using two technical replicates of each of three biological replicates per cell condition (n=6). Labels follow B. The result was significantly different by T-test (p<0.001) after removal of one Scr outlier (panel displayed) or by Mann-Whitney U test (p<0.002) including the outlier. (D) SAM ion intensities in the cells from B were not significantly different (T-test). Labels follow B. (E) 1-MNA attenuates rounded morphology in DM cells. Results for n=5 for each 1-MNA concentration are shown in the boxplot. 0.5 mM 1-MNA was significantly different (p<0.0001, ANOVA, post hoc Tukey HSD) to all other treatments.

**Figure S5.**
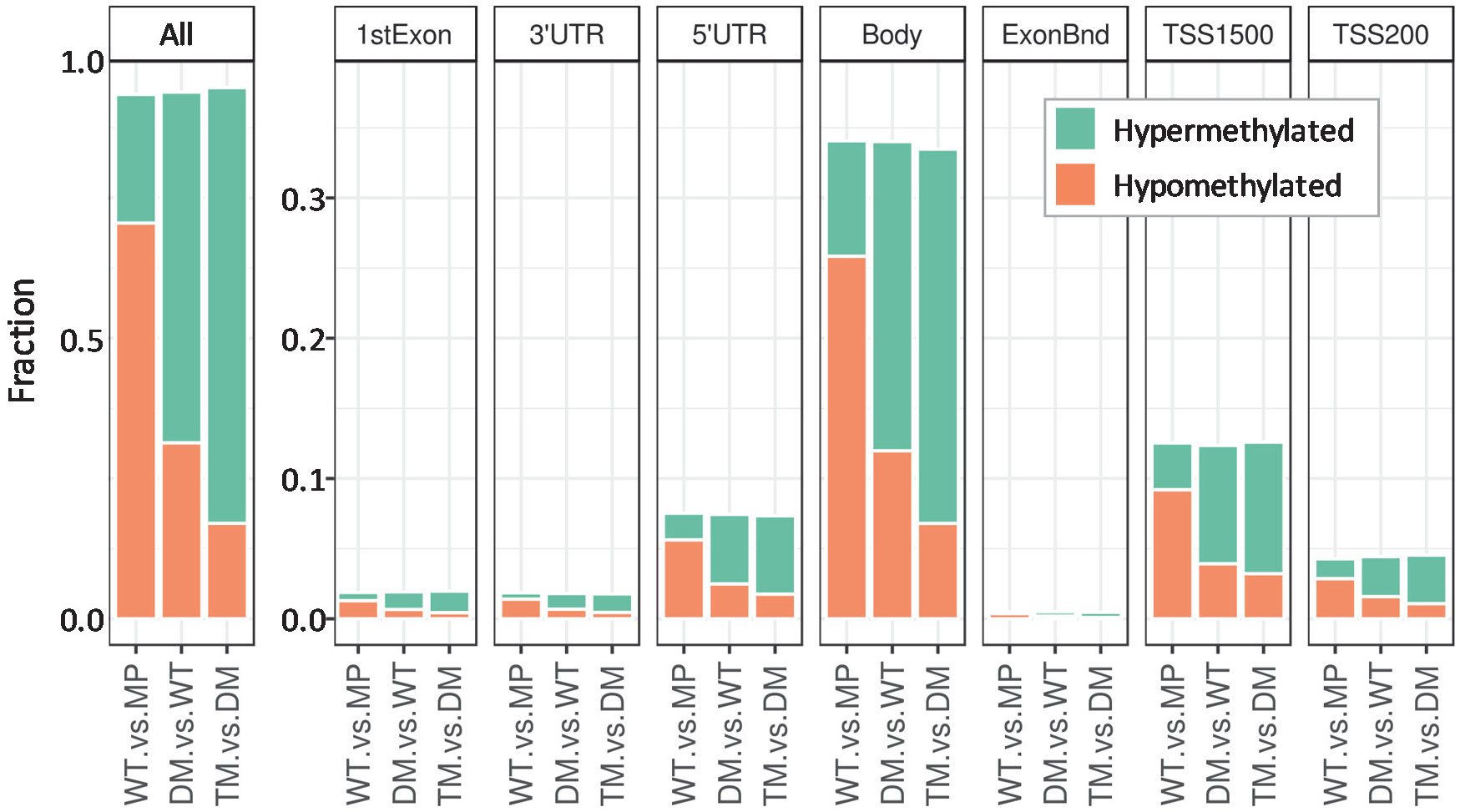
Methylation status of CpG associated with annotated coding genes. Related to Figure 7. Significant differentially methylated probes were annotated with the UCSC gene feature and the proportion of probes attributed to each feature counted. The y-axis shows the frequency of hypermethylated-hypomethylated probes per feature, and the x-axis shows the difference in those frequencies between WT/MP, DM/WT and TM/DM. 3’UTR: 3’ untranslated region; 5’UTR: 5’ untranslated region; Body: Between the ATG and stop codon; irrespective of the presence of introns, exons, TSS, or promoters; ExonBnd: exon boundaries: TSS1500/TSS200: within indicated number of residues of transcription start site (TSS).

**Figure S6.**
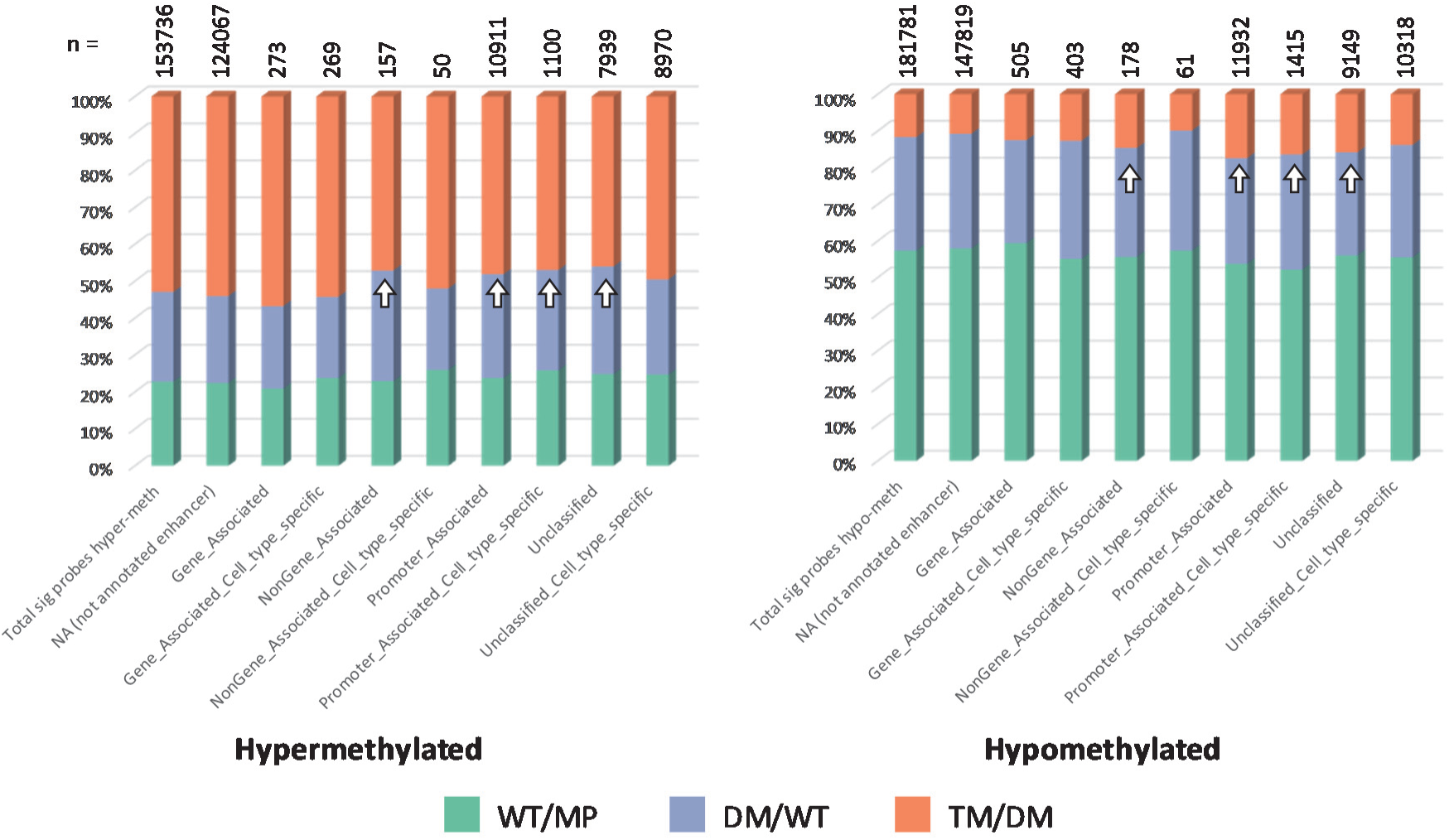
Methylation status of CpG associated with annotated enhancers. Related to Figure 7. Significant differentially methylated probes were annotated with the 450k_Enhancer feature and the proportion of probes attributed to each feature counted. Arrows indicate increased hypomethylation and reduced hypermethylation of enhancers in the TM/DM comparison.

**Figure S7.**
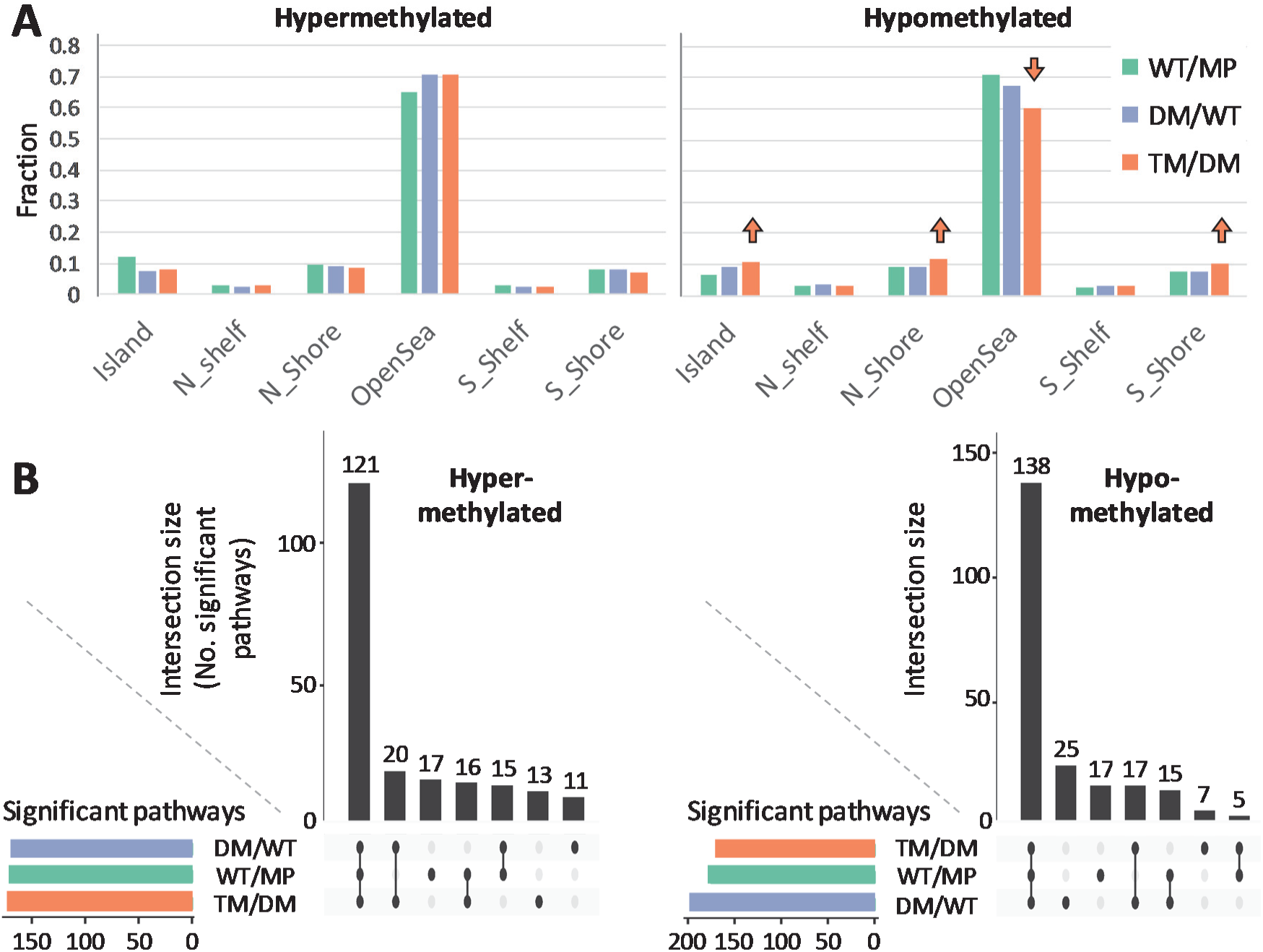
Chromosomal context of differentially methylated CpG, and biological association of genes. Related to Figure 7. (A) CpG feature classifications for differentially methylated probes. The percentage of all significant probes that were hypermethylated or hypomethylated per CpG feature is presented for each comparison. Up arrows point to relatively greater fractions of Island and Shore genomic regions for the TM/DM comparison. The down arrow points to reduced OpenSea hypomethylation in the TM/DM comparison. (B) Intersections of significant KEGG pathways between WT/MP, DM/WT and TM/DM. KEGG enrichment was applied separately to significant hypermethylated and hypomethylated probes. The main bar chart presents the number of pathways common to each comparison. Each bar represents an intersect notated by linked dots on the x-axis. The number of significant pathways per comparison is presented as a horizontal bar chart. The KEGG results are in File S1 and the list of pathways unique to each comparison are provided. Pathways analysis of probes genes corresponding to the top 2000 most variable differentially methylated probes in CpG Island or Shore chromosomal regions. Probes were separated into hypermethylated and hypomethylated data sets from D, including only probes from Island or Shores. Enriched Gene Ontology (GO) pathways enrichments are shown. The identities of pathways uniquely differential to one of the three cell type comparisons (WT/MP, DM/WT, or TM/DM) for both hypo- and hyper-methylated data sets and KEGG pathways are available in Supplemental Information File 1.

**Table S1.**
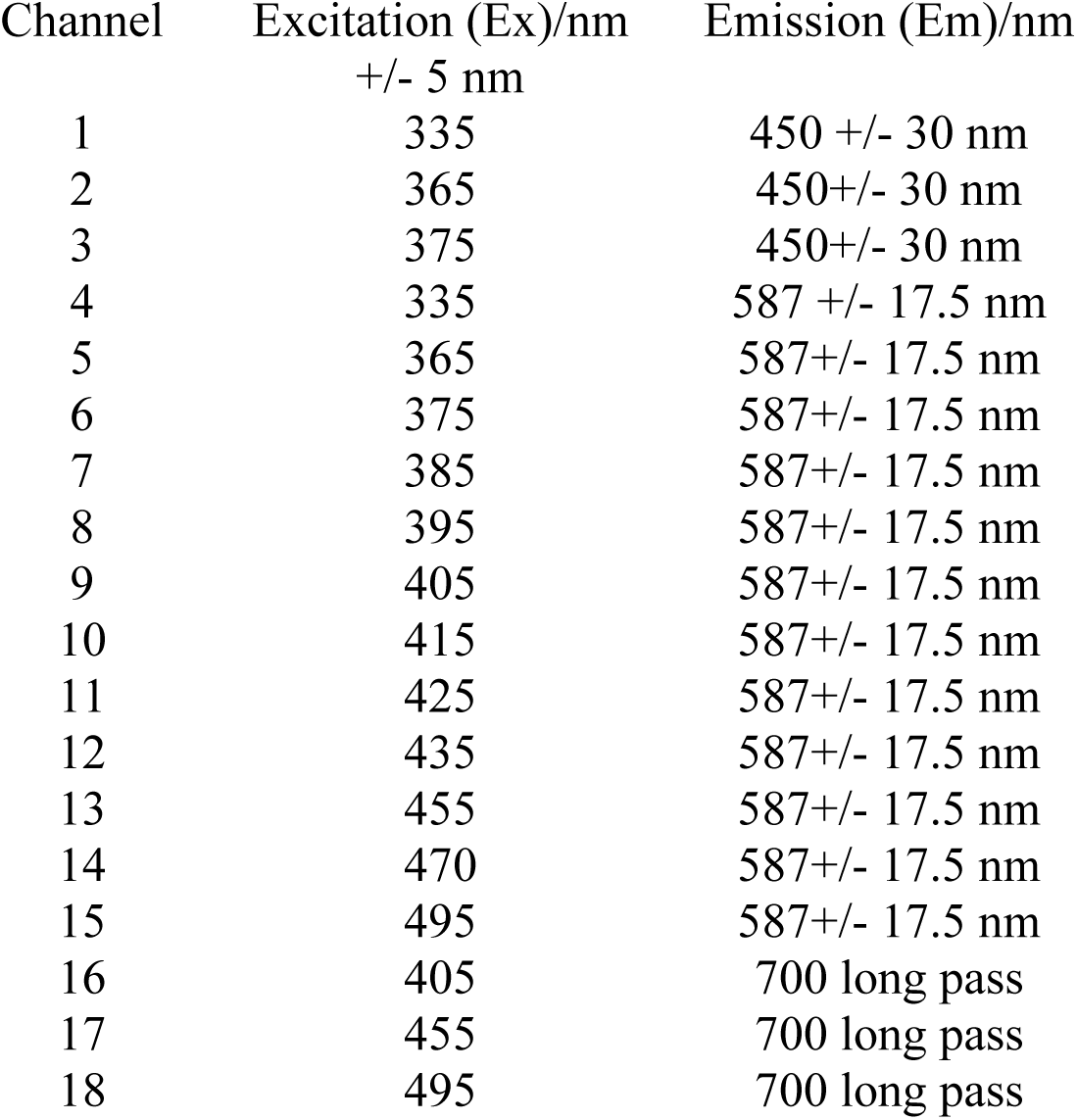
Spectral channel filters employed for hyperspectral autofluorescence imaging. Related to Figure 1. For further details of this approach see Gosnell et al. (Gosnell et al., 2016a).

**Table S2.** GO pathways enrichment results. Related to Figure 7. The top ten GO enrichments for each cell comparison from Figure 7. Full results are available in File S1.

| Comparison | GO_ID | FDR | PValue | sigGO.Term |
| --- | --- | --- | --- | --- |
| MP/WT | GO:0032501 | 8.1E-07 | 1.85E-10 | multicellular organismal process |
|  | GO:0048731 | 2.03E-06 | 9.28E-10 | system development |
|  | GO:0007275 | 5.75E-05 | 4.09E-08 | multicellular organism development |
|  | GO:0030154 | 5.75E-05 | 6.05E-08 | cell differentiation |
|  | GO:0048869 | 5.75E-05 | 6.58E-08 | cellular developmental process |
|  | GO:0048856 | 0.00013 | 1.79E-07 | anatomical structure development |
|  | GO:0022610 | 0.000141 | 2.26E-07 | biological adhesion |
|  | GO:0032502 | 0.00017 | 3.52E-07 | developmental process |
|  | GO:0023052 | 0.00017 | 3.67E-07 | signaling |
|  | GO:0009653 | 0.00017 | 3.89E-07 | anatomical structure morphogenesis |
| WT/DM | GO:0032501 | 1.79E-07 | 4.14E-11 | multicellular organismal process |
|  | GO:0009653 | 2.27E-05 | 1.05E-08 | anatomical structure morphogenesis |
|  | GO:0048731 | 5.71E-05 | 4.5E-08 | system development |
|  | GO:0050896 | 5.71E-05 | 5.28E-08 | response to stimulus |
|  | GO:0023052 | 6.44E-05 | 7.44E-08 | signaling |
|  | GO:0007165 | 0.000153 | 2.12E-07 | signal transduction |
|  | GO:0007154 | 0.000227 | 3.7E-07 | cell communication |
|  | GO:0051239 | 0.000227 | 4.2E-07 | regulation of multicellular organismal process |
|  | GO:0051716 | 0.000241 | 5.01E-07 | cellular response to stimulus |
|  | GO:0007275 | 0.000325 | 7.51E-07 | multicellular organism development |
| TM/DM | GO:0032501 | 2.71E-07 | 6.56E-11 | multicellular organismal process |
|  | GO:0048731 | 4.52E-05 | 2.19E-08 | system development |
|  | GO:0009653 | 5.33E-05 | 3.91E-08 | anatomical structure morphogenesis |
|  | GO:0007275 | 5.33E-05 | 5.16E-08 | multicellular organism development |
|  | GO:0007399 | 0.000113 | 1.37E-07 | nervous system development |
|  | GO:0050896 | 0.000141 | 2.16E-07 | response to stimulus |
|  | GO:0032502 | 0.000141 | 2.39E-07 | developmental process |
|  | GO:0048856 | 0.000173 | 3.36E-07 | anatomical structure development |
|  | GO:0030154 | 0.000214 | 4.67E-07 | cell differentiation |
|  | GO:0048869 | 0.000365 | 8.84E-07 | cellular developmental process |

**Table S3.**
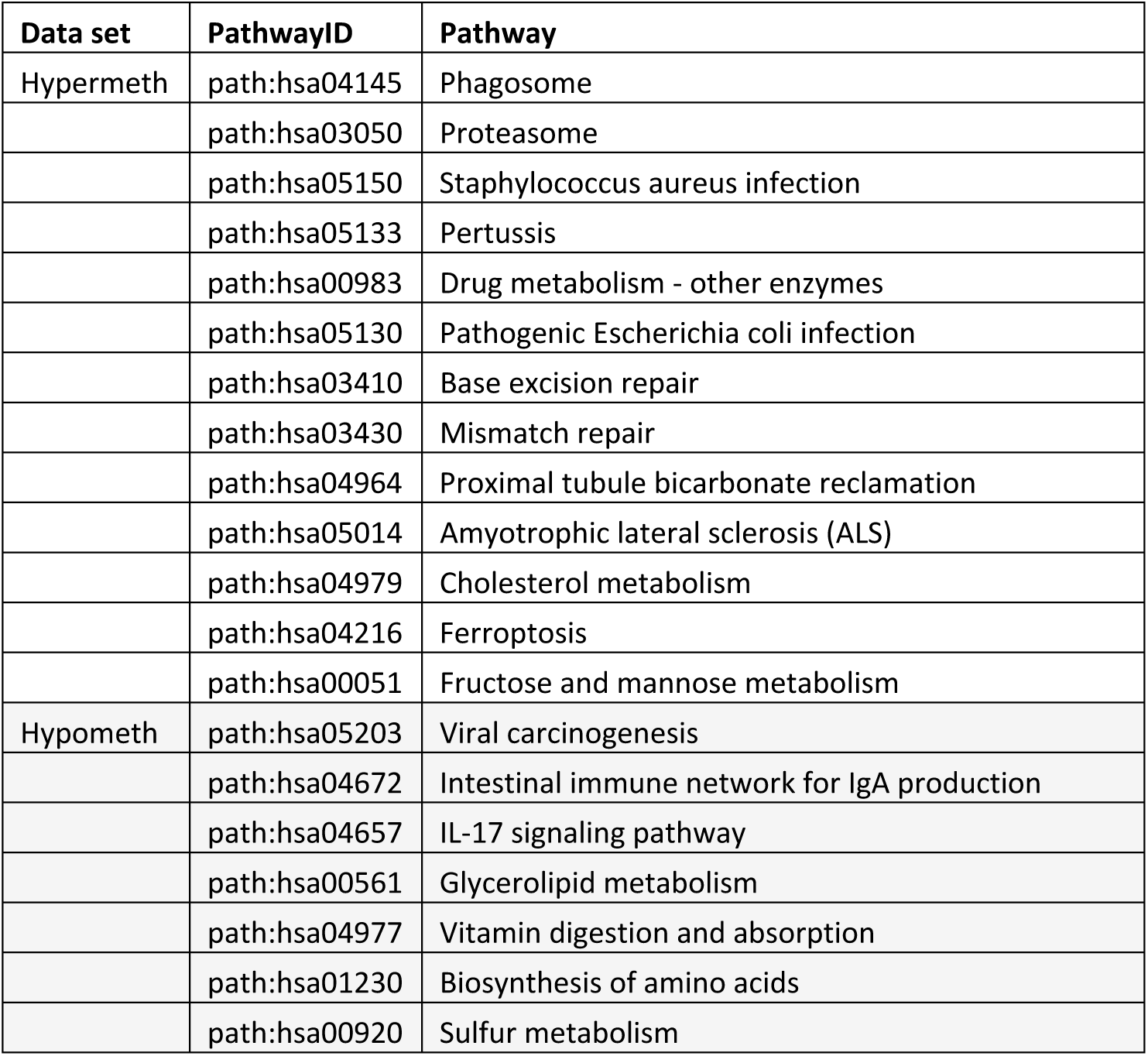
Unique KEGG pathways detected for TM/DM hyper and hypo-methylated gene sets. Related to Figure 7. The identities of the 11 hypermethylated and 7 hypomethylated pathways unique to the TM/DM comparison from Figure S7B are given. Full results are available in File S1.

### Supporting Information file

**File S1**. A zip archive containing excel files with the most significant pathways enrichment results from methylomics analysis including those of Figure S7B, Table S2 and Table S3. The archive unpacks as five separate folders, each containing the following pathways analysis results.

A) All GO enrichments for each cell comparison significant below the adjP =0.001 level.

B) All GO enrichments that were uique to a particular cell comparison.

C) KEGG enriched pathways for each cell comparison, with analyses performed separately on hypermethyalted (files labelled up) and hyperemethylated (files labelled down) probes. These correspond to the pathways of of Figure S7B.

D) All KEGG enriched pathways from C which were unique to a particular cell comparison.

E) All All Reactome enriched pathways for each cell comparison significant below the adjP =0.05 level.

